# PM_2.5_ Exposure Facilitates SARS-CoV-2 Infection through ACE2/TMPRSS2 Regulation and Suppression of Anti-Viral Response

**DOI:** 10.1101/2025.10.02.679237

**Authors:** Hadi Rajabi, Ozgecan Kayalar, Gulen Esken, Fusun Can, Hasan Bayram

## Abstract

**Background:** Epidemiological studies suggest an interaction between air pollution including particulate matter <2.5 µm (PM_2.5_) and coronavirus disease 2019 (COVID-19) mortality and morbidity; however, the underlying mechanisms are not clear. The aim of our study was to investigate effects of PM_2.5_ on viability, epithelial integrity, and cellular entry of SARS-CoV-2 into airway epithelial cells, and the mechanisms involved.

**Methods:** We exposed Calu-3 airway epithelial cell cultures to PM_2.5_ (10, 50, and100 µg/ml) and SARS-CoV-2 (MOI 0.01) for 24 h. The viability of Calu-3 cells and epithelial barrier integrity were determined using MTT assay and immunofluorescence staining for Zonula Occludens-1, respectively. mRNA expression for viral entry-related genes such as angiotensin converting enzyme (*ACE)2* and transmembrane protease, serine (*TMPRSS)2,* and inflammatory and inflammasomal genes, including interleukin (IL)-8,IL-6, nuclear factor (NF)-κB p65 (*RELA*), *JNK, c-JUN, Caspase-1, IL-1*β, *NLRP3,* was analyzed by qRT-PCR. Intracellular viral spike protein intensity and RNA-dependent RNA polymerase (RdRP) expression were determined using immunofluorescence staining and qRT-PCR, respectively. ELISA was used to analyze the release of inflammatory cytokines (IL-8, IL-6, and GM-CSF).

**Results:** Higher concentrations of 100µg/ml PM_2.5_ decreased Calu-3 cell viability (p=0.02) and deteriorated epithelial barrier integrity, while 50 µg/ml of PM_2.5_ (p<0.01) induced mRNA expression for *ACE2* and *TMPRSS2.* Although PM2.5 alone decreased c-JUN, it did not alter the expression of mRNA for JNK and RELA. In contrast, a combination of SARS-CoV-2 and PM_2.5_ led to a significant increase in mRNA for both JNK and RELA (p < 0.05 and p < 0.01, respectively) and attenuated c-JUN expression. Moreover, our results indicated an increase in the expression of IL-1β, IL-6, and GM-CSF following exposure to PM_2.5_ and PM_2.5_ + SARS-CoV-2, whereas IL-8 was induced only by SARS-CoV-2 exposure. Co-incubation of Calu-3 cells with PM_2.5_ and SARS-CoV-2 leads to a decrease in IL-8, IL-1β, *Caspase-1 (CASP-1),* and Interferon gamma *(IFNG*) expression. Finally, the viral load (RdRP) also increased in the presence of both PM_2.5_ and the SARS-CoV-2 group.

**Conclusion:** Our findings have demonstrated that PM_2.5_ impaired epithelial integrity and cell viability, whereas it increased the mRNA expression for *ACE2* and *TMPRSS2,* and induced inflammatory changes in Calu-3 cells incubated with SARS-CoV-2. These findings suggest that PM_2.5_ can facilitate the entry of SARS-CoV-2 into airway epithelial cells, and that both PM_2.5_ and SARS-CoV-2 can decrease the inflammatory and antiviral responses of the host cell.

## 1. Introduction

Coronavirus disease 2019 (COVID-19), which is induced by severe acute respiratory syndrome coronavirus 2 (SARSl7lCoVl7l2), one of the β-type coronaviruses, has affected almost all world population, since its onset in 2019, and the struggle against the infection is still ongoing because new variants exist [1, 2]. The severity of COVID-19 has been shown to depend on different factors, ranging from genetic and metabolic disorders to environmental factors [3, 4].

Studies suggest that individual susceptibility and environmental conditions, such as outdoor air pollution, can contribute to the transmission of SARS-CoV-2 and the mortality and morbidity caused by COVID-19 [5–8]. An increase of 1 μg/m^3^ PM_2.5_ was associated with increased COVID-19 deaths [9]. Similarly, Semczuk-Kaczmarek et al. found that an annual average increase of PM_2.5_ by approximately 10 μg/m³ led to a 36.7% rise in COVID-19 cases and a 29% increase in the mortality rate[10]. Furthermore, Zhang et al. reported that each 0.5 µg/m³ increase in PM_2.5_ concentration corresponded to a 10% rise in COVID-19 cases, 9% increase in COVID-19-related hospitalizations, and 23% rise in mortality [11, 12].

SARS-CoV-2 primarily enters cells through the receptors for angiotensin-converting enzyme 2 (ACE2) and transmembrane protease, serine 2 (TMPRSS2) [13]. Because of their small size, PM_2.5_ can reach the lower parts of the respiratory system, including alveoli, and cause epithelial damage, inflammation, and oxidative stress [14]. Research has shown that PM_2.5_ can enhance SARS-CoV-2 viral entry by inducing the expression of these receptors. A recent study by Batto et al. reported that PM_2.5_ increased the expression of ACE2 by 40% in a mouse model [15]. Studies also suggest that PM_2.5_ may play a role in the transmission and dissemination of SARS-CoV-2 [16, 17]. Indeed, PM collected from hospital gardens and various urban locations were found to contain viral particles such as RNA-dependent RNA polymerase (RdRP) and the Nucleocapsid (NP) N1 fragment [16, 18].

The innate immune system is the first defense against viral infection, triggering antiviral mediators that restrict viral replication and activate immune cells [19]. The inflammasomal response plays a critical role in microbial and viral infection by inducing an inflammatory reaction. However, the interaction between PM_2.5_ and SARS-CoV-2 at the cellular level is not clear, and there are no specific studies about the innate immune system and inflammasomal response in SARS-CoV-2 in the presence of PM_2.5._

In this study, we investigated the interaction between PM_2.5_ and SARS-CoV-2 and found that PM_2.5_ could induce cellular entry of the virus by increasing the expression of ACE2 and TMPRSS2. Furthermore, PM_2.5_ altered the inflammasomal response in respiratory epithelial cells infected by SARS-CoV-2.

## 2. Methods

### 2.1. Extraction and isolation of particulate matter ≤ 2.5 µm (PM_2.5_)

PM_2.5_ samples were collected at the air quality monitoring station in the Alibeyköy, Istanbul in cooperation with the Istanbul Metropolitan Municipality (IMM) at a flow rate of 300 m^3^/min for 24 h on 3-5 cm diameter polytetrafluoroethylene polymer (PTFE) membrane filters (Millipore, USA) using a high-volume air sampler with adapter (Anderson Instrument Co. USA). After sampling, the filters were divided into two equal parts. One half will be split into smaller pieces and overwhelmed in ultrapure water (18.2 ohms), and the particulate matter passed into the solution by ultrasonication for about 30 min at a frequency of 60kHz. The suspension was filtered through a 40 µm pore diameter injector filter (Sarstedt AG, Germany) to remove fiber fragments. Then, the suspension was dried by vacuum lyophilizer and sterilized using ethylene oxide. Then, the PMs were diluted in Dulbecco’s Phosphate-Buffered Saline (DPBS,1X, Biowest, France) with a ratio of 1:1 and stored at −20_JC for further experiments.

### 2.2. Calu-3 Cell culture and PM _2.5_ and SARS-CoV-2 treatment

Calu-3 cells were cultured in Eagle’s Minimum Essential Medium (EMEM) (Wisent, USA) supplied with 10% foetal bovine serum (FBS) (Gibco, Germany), 1% sodium pyruvate (Gibco, UK) and 1% penicillin-streptomycin (Biowest, France) at 75 cm^2^ cell culture flask in humid condition with 5% CO_2_ and 37_J C. After reaching 80% confluency, cell cultures were subcultured using trypsin/EDTA (0.25% w/w; Wisent, USA). The culture medium was changed every 48 h. For PM_2.5_ treatment Calu-3 cells were incubated in serum-free media (SF) for 24 h, subsequently, cells were exposed to three concentrations of PM_2.5_ (0,10, 50, and 100 µg/ml) for 24 h. The viral treatment of Calu-3 cells was performed in the Biosafety Level 3 (BSL-3) laboratory at the Koç University İşBank Center for Infectious Diseases (KUISCID). Before infection, the cell cultures were incubated in SF media for 24 h and then treated with PM_2.5_ for 24 h before SARS-CoV-2 viral infection. The used dosage of the virus was 0.01 multiplicity of infection (MOI=0.01) for 1 h, after finishing the viral infection medium was refreshed with fresh SF medium and incubated for an additional 24h.

### 2.3. Assessment of Calu-3 Cell Viability

To investigate the effective concentration of PM_2.5_ on Calu-3 cell viability, the staining method based on 3-(4,5-dimethylthiazol-2-yl) 2,5-diphenyltetrazolium bromide (MTT) uptake by living cells and reduction by mitochondrial oxidoreductase enzymes was used (Active form). The cells were cultured in 24 well plates and incubated for 24 h in EMEM serum-free medium prior to incubation with three different concentrations of PM_2.5_ (0,10, 50 and 100 µg/ml). Following 24 h of PM_2.5_ exposure, the supernatant was collected and replaced with 500µl of MTT solution and incubated for 90min. Then, the MTT solution was discarded and 500 µl of dimethylsulfoxide (DMSO; PanReac, Germany) was added to each well to dissolve the formazan crystals and mixed for 5 min at 100-150 rpm. Subsequently, the colour change was evaluated using a microplate reader (Biotek, USA) at a wavelength of 540 nm [20].

### 2.4. Analysis of Calu-3 cells’ barrier function using immunofluorescence staining and live cell imaging

To determine the effect of PM_2.5_ on cell culture barrier function, 10^4^ Calu-3 cells were cultured in 96-well plates and treated with PM_2.5_. Cells were then fixed with 4% paraformaldehyde for 20 min at room temperature, and after washing with DPBS, permeabilization was done with 0.1% Triton X-100. After blocking with superblock, cells were incubated with Zonula Occludens-1 (ZO-1) rabbit polyclonal primary antibody (AF5145, Affinity Biosciences) overnight at +4°C and then incubated with Alexa Fluor 594 labeled goat anti-rabbit secondary antibody (Invitrogen, USA) for 90 min in 37°C. After finishing the staining, the cells were visualized using a fluorescence microscope (IX71; Olympus, Japan). The images were analyzed using ImageJ 1.52e (NIH, MD, USA) [21].

### 2.5. Analysis of SARS-CoV-2 infection in Calu-3 cells using immunofluorescence staining and live cell imaging

To determine the effect of various concentrations of PM_2.5_ on viral entry rate, immunofluorescence staining (IF) of viral spike protein (S protein) was done. For this goal, 10^4^ Calu-3 cells were cultured in 96 well plates and treated with PM _2.5_ and SARS-CoV-2, cells were then fixed with 4% paraformaldehyde for 20 min at room temperature and after washing with PBS, permeabilization has done with 0.1% Triton X-100. After blocking with superblock, cells were incubated with viral spike protein-specific mouse primary antibody (ab273433, Abcam, UK) overnight at +4°C and then incubated with Alexa Fluor 488 labeled goat-anti mouse secondary antibody (Invitrogen, USA) for 90 min in 37°C. After finishing the staining, the cells were visualized using a fluorescence microscope (IX71; Olympus, Japan).

### 2.6. Bioinformatic analysis of possible genes and pathways

For the exact selection of genes and pathways involved in innate immune system modification in the presence of PM_2.5_ and SARS-CoV-2, we used the Genemania.org database and selected more relevant and related genes[22].

### 2.7. RNA isolation and Real-Time PCR

Cells seeded at a density of 3×10^5^ in a 6-well plate culture dish and incubated with PM_2.5_ in the presence and absence of SARS-CoV-2. Infected cells were neutralized and collected by lysis buffer and Trizol and total RNA isolation was performed with an RNA isolation kit (Zymo, USA). After measuring the RNA quantity and purity with a Nanodrop^TM^ 2000c (ThermoFisher Scientific, MA, USA) device, cDNA was synthesized from RNA by using an iScript cDNA synthesis kit (Biorad, USA). Expression of inflammatory and viral-related genes such as *ACE2*, *TMPRSS2*, MAPK8 (*JNK)*, *NF-*κ*B p65 (*RELA), the *NOD-like receptor protein (NLRP)3, CASP-1,* IL-1β, *IL-8 (CXCL-8), and Interferon gamma (IFNG)* was analyzed by the QuantStudio^TM^ 7 qRT-PCR instrument (Applied Biosystems, USA) and normalized to GAPDH, which was used as an internal housekeeping gene. The fold increase of gene expression between groups was calculated using the 2^-ΔΔCt^ method [23]. The primer sequences are outlined in supplementary Table 1. Also, to assess SARS-CoV-2 RdRP expression, RNA isolation was performed from collected supernatants using (Qiagen, Germany). cDNA was synthesized from RNAs by using an iScript cDNA synthesis kit (Biorad, USA). The viral qRT-PCR was performed using TaqMan™ Fast Advanced Master Mix and FAM probe-based QuantStudio 7 qRT-PCR instrument (Applied Biosystems, USA).

### 2.8. Enzyme-Linked Immunosorbent Assay (ELISA)

Concentrations of pro-inflammatory cytokines such as IL-6, IL-8, and GM-CSF in the infected Calu-3 cell culture supernatant were measured using enzyme-linked immunosorbent assay (ELISA) kit (R&D Systems, Minneapolis, MN). according to the manufacturer’s instructions.

### 2.9. Statistical Analysis

All experiments were performed with appropriate controls in duplicate or triplicate. Data were tested for normality using the D’Agostino-Pearson Test. One-way analysis of variance (ANOVA) or Kruskal-Wallis tests were performed, and appropriate post-hoc tests were applied accordingly. Comparisons between two groups were made using the unpaired t-test or the Mann-Whitney U test. Correlation analyses were performed using Spearman correlation. P values less than 0.05 were considered significant. Statistical analysis was performed using Prism V.6.07 (GraphPad).

## 3. Results

### 3.1. Effects of PM_2.5_ on Calu-3 cell viability

After treating Calu-3 cells with three different concentrations of PM_2.5_ (0,10, 50, and 100 µg/ml) for 24h, we found that only 100 µg/ml of PM_2.5_ (median= 24, Q1=34.39, and Q3=58.86 nm) significantly decreased cell viability as compared to 0µg/ml PM_2.5_ (median= 112.8, Q1=108, and Q3=128 nm, p<0.02) (**Figure 1**).

**Figure 1.**
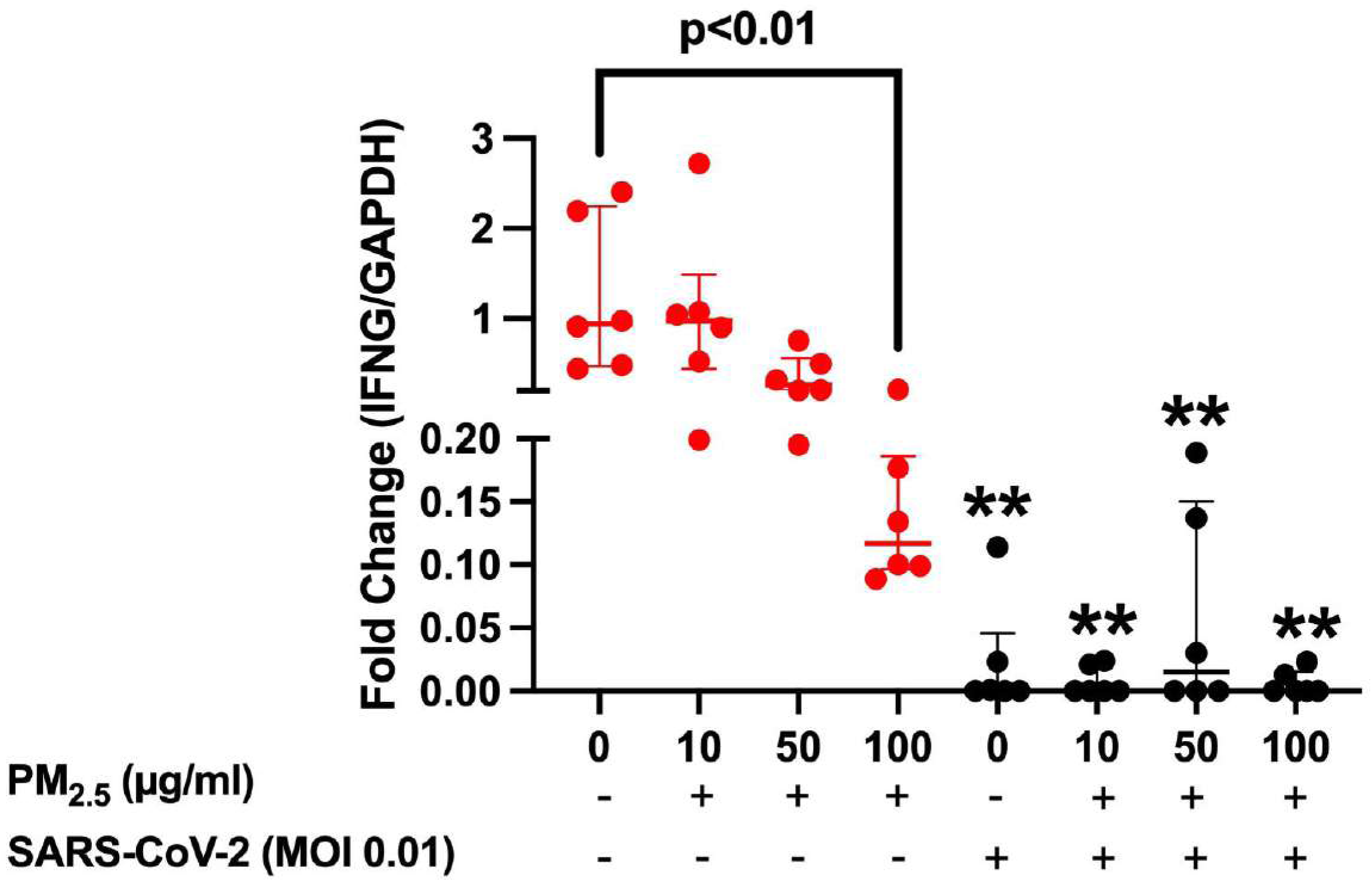
Effects of 0,10, 50, and 100 µg/ml particle matter ≤ 2.5µm (PM_2.5_) on Calu-3 cell viability.

### 3.2. **Effect of** PM_2.5_ on Calu-3 cell culture barrier integrity

IF staining for ZO-1 showed decreased barrier integrity of Calu-3 cells incubated with 100 µg/ml PM_2.5_ compared to 0 µg/ml PM_2.5_ group (p=0.0015). Other concentrations of PM_2.5_ used (10 and 50 µg/ml) were ineffective (**Figure 2**).

**Figure 2.**
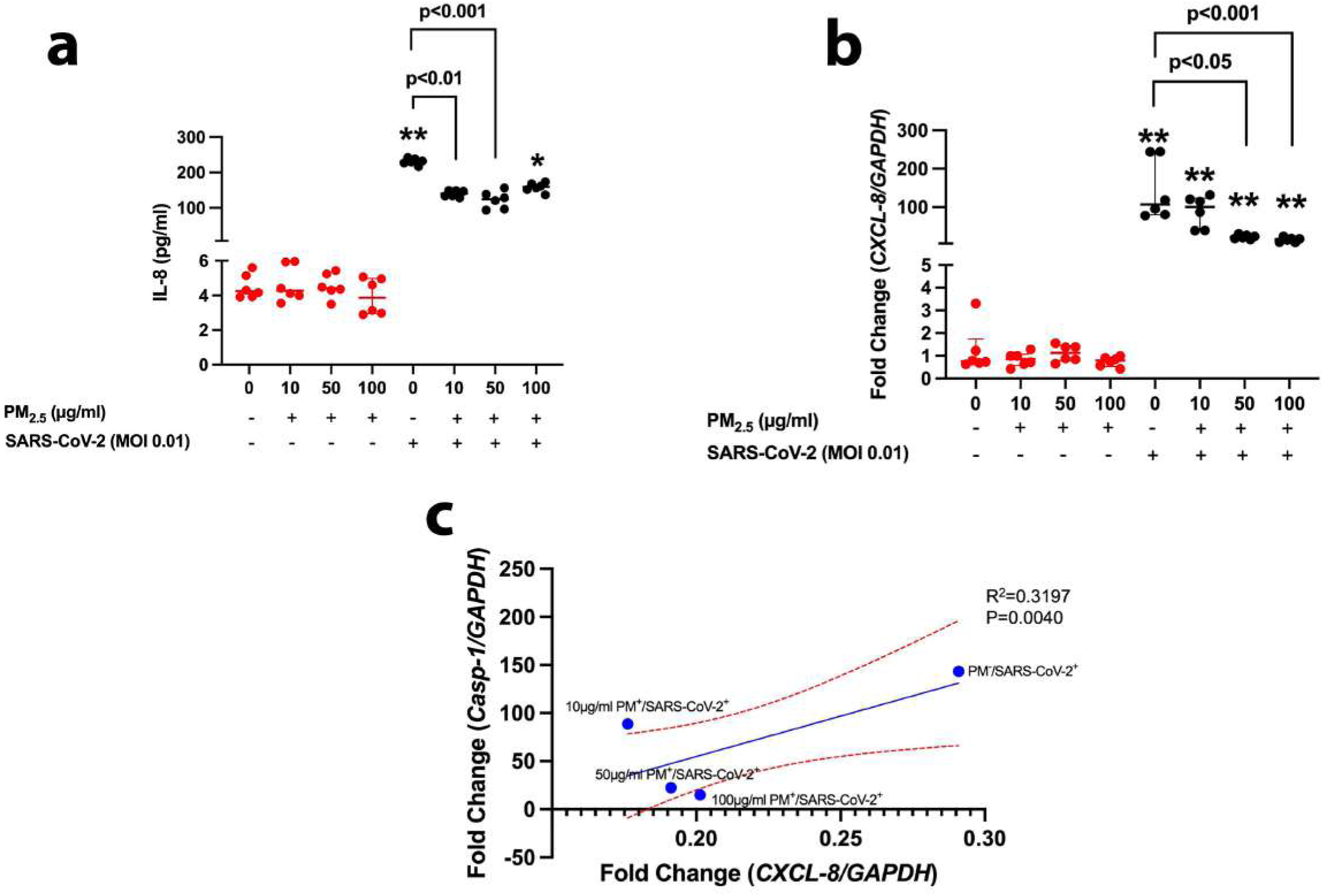
Effects of 0,10, 50, and 100 µg/ml particle matter ≤ 2.5µm (PM2.5) on ZO-1 junctions in Calu-3 cell cultures. (Magnification=20x).

### 3.3 The fluorescence intensity of the viral spike protein and RdRP gene expression in Calu-3 cells exposed to PM_2.5_

Viral S protein immunofluorescence staining of Calu-3 cells treated with PM_2.5_ and SARS-CoV-2 showed stronger staining for intracellular S protein following increasing concentration of PM_2.5_ (**Figure 3a**). Also, Real-Time PCR results have demonstrated that RdRP gene expression was increased by 50 µg/ml PM_2.5_ (p<0.01) (**Figure 3b**).

**Figure 3.**
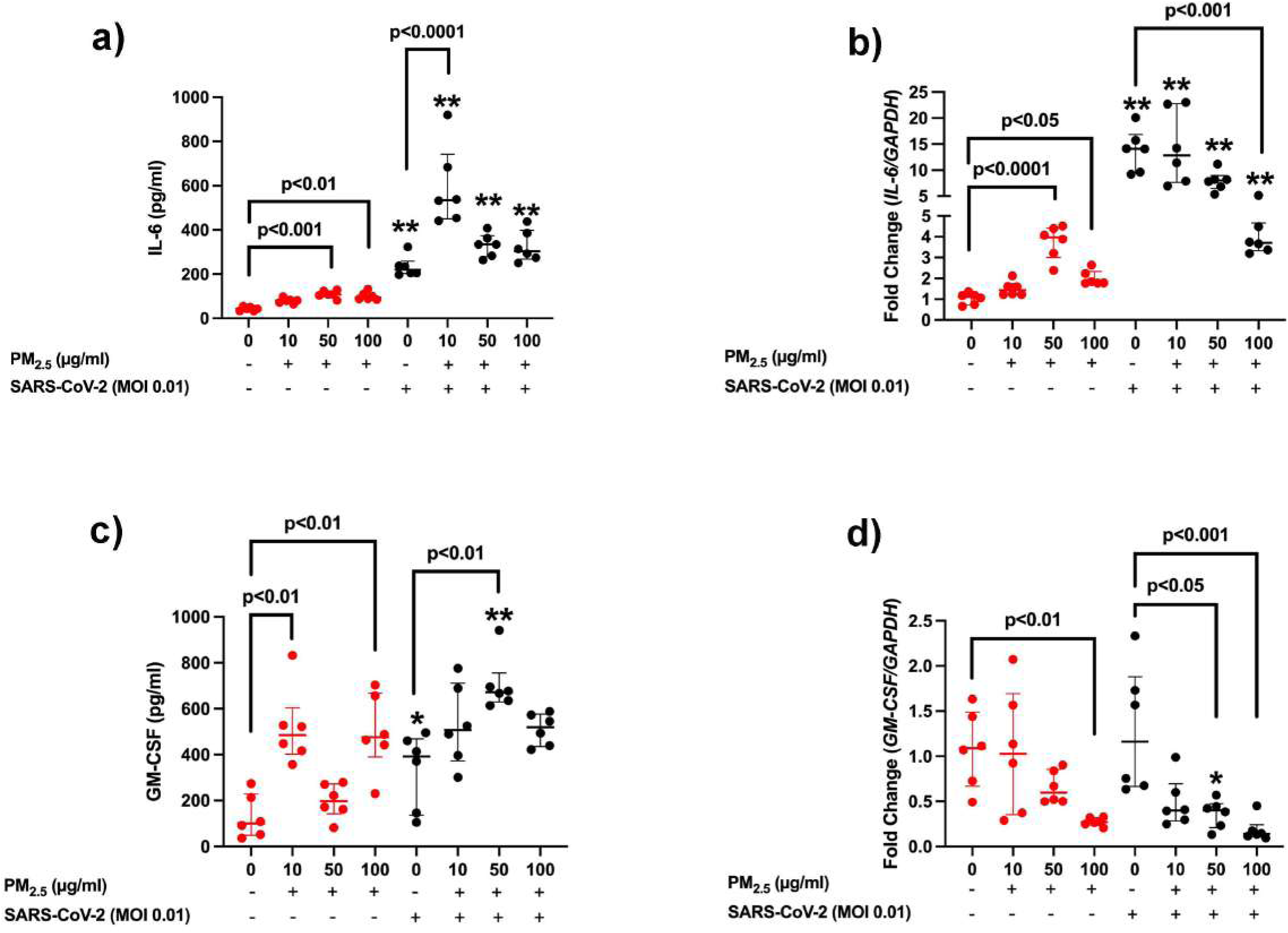
(a) Representative immunofluorescence images of intracellular SARS-CoV-2 spike (S) protein in Calu-3 cells exposed to 0-<u>100</u>µ<u>g/ml</u> of particle matter ≤ 2.5µm (PM_2.5_) (Magnification=10x). (b) Effects of 50 µg/m particle matter ≤ 2.5µm (PM_2.5_) and SARS-CoV-2 (MOI=0.01) on mRNA expression for RdRP.

### 3.4. Effects of co-incubation of PM_2.5_ and SARS-CoV-2 on mRNA expression for ACE2 and TMPRSS2 in Calu-3 cells

To understand the effect of PM_2.5_ on SARS-CoV-2 viral entry genes in Calu-3 cells, we evaluated *ACE2* and *TMPRSS2* gene expression cells incubated with different concentrations of PM_2.5_ in the absence and presence of SARS-CoV-2. Based on our results, gene expression for both *ACE2* and *TMPRSS2* was increased by 50 µg/ml PM_2.5_ (p<0.01) (**Figures 4a and b**). Similarly, SARS-CoV-2 itself increased *ACE2* mRNA expression compared to the untreated control group (p<0.05) (Figure 4a). However, co-treatment of Calu-3 cells with PM_2.5_ (50 µg/ml) and SARS-CoV-2 led to a decreased expression of *ACE2* mRNA (p<0.05), whereas *TMPRSS2* mRNA expression was increased by 100 µg/ml PM_2.5_ (p<0.001, **Figures 4a and b**).

**Figure 4.**
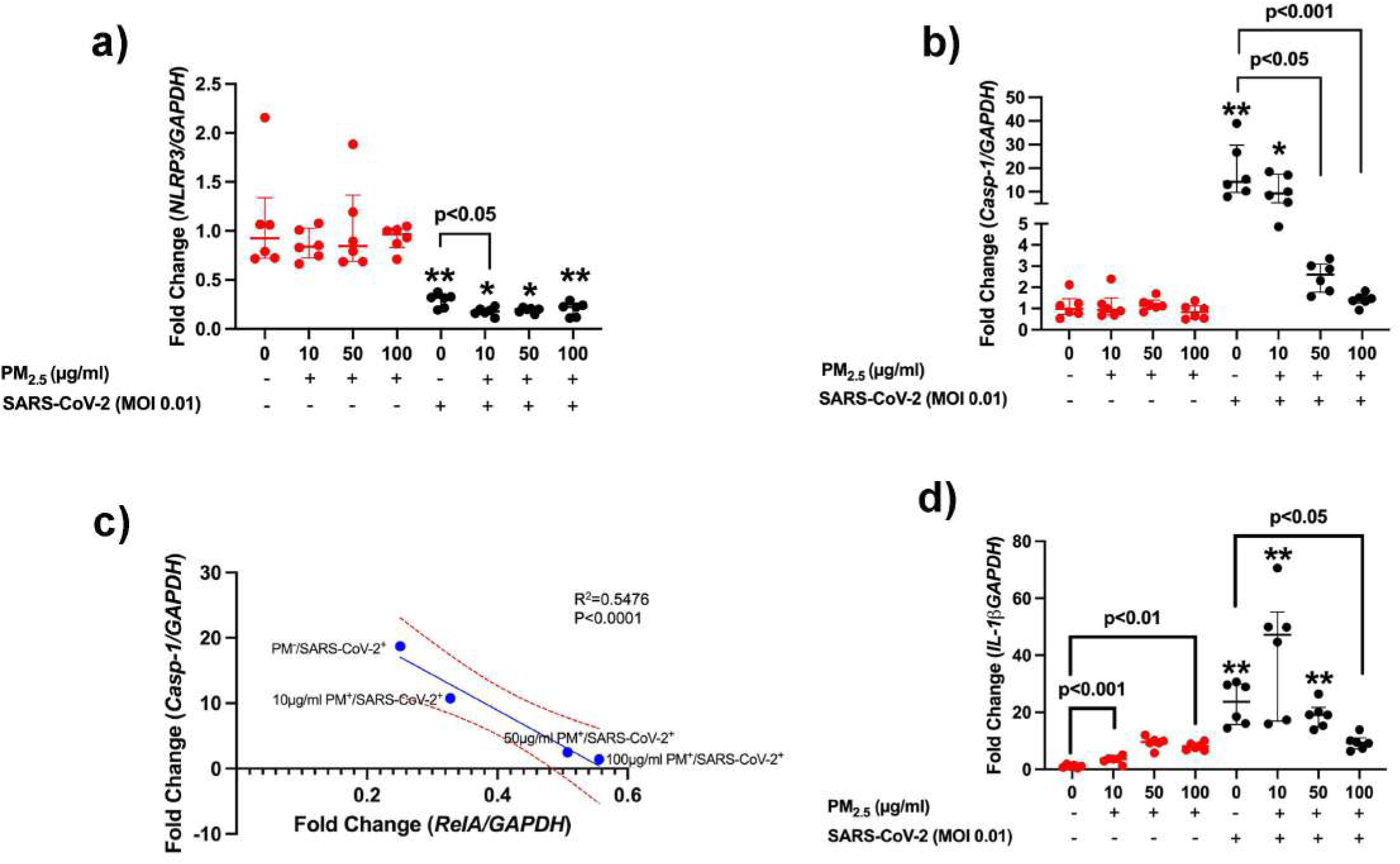
Effects of particle matter ≤ 2.5µm (PM_2.5_) (0-100µg/m) and SARS-CoV-2 (MOI=0.01) on mRNA expression for (a) ACE2, and (b) TMPRSS2 in Calu 3 cells following 24 hours incubation (*p<0.05 vs corresponding PM_2.5_ concentrations without SARS-CoV-2).

### 3.5. Effects of PM_2.5_ and SARS-CoV-2 on the mRNA expression for JNK, JUN, and NF-**κ**B p65 (RELA) in Calu-3 cells

The gene expression analyses showed that although PM_2.5_ alone did not alter the JNK mRNA expression, its expression was significantly increased by 50 (p<0.05) and 100 (p<0.0001) µg/ml PM_2.5_ in the presence of SARS-CoV-2, as compared to SARS-CoV-2 alone. Furthermore, when comparing with the corresponding PM_2.5_ concentrations, we found that in the presence of SARS-CoV-2, the JNK mRNA expression induced by 10 (p<0.05) and 100 (p<0.01) µg/ml PM_2.5_ was significantly higher (**Figure 5a**). However, 100 µg/ml PM_2.5_ (p<0.05) significantly reduced JUN mRNA expression compared to 0µg/ml PM_2.5_. Similarly, SARS-CoV-2 (p<0.001) alone, together with the presence of 10 (p<0.001) and 50 (p<0.01) µg/ml PM_2.5,_ led to a significant decrease in *JUN* expression. (**Figure 5b**).

**Figure 5.**
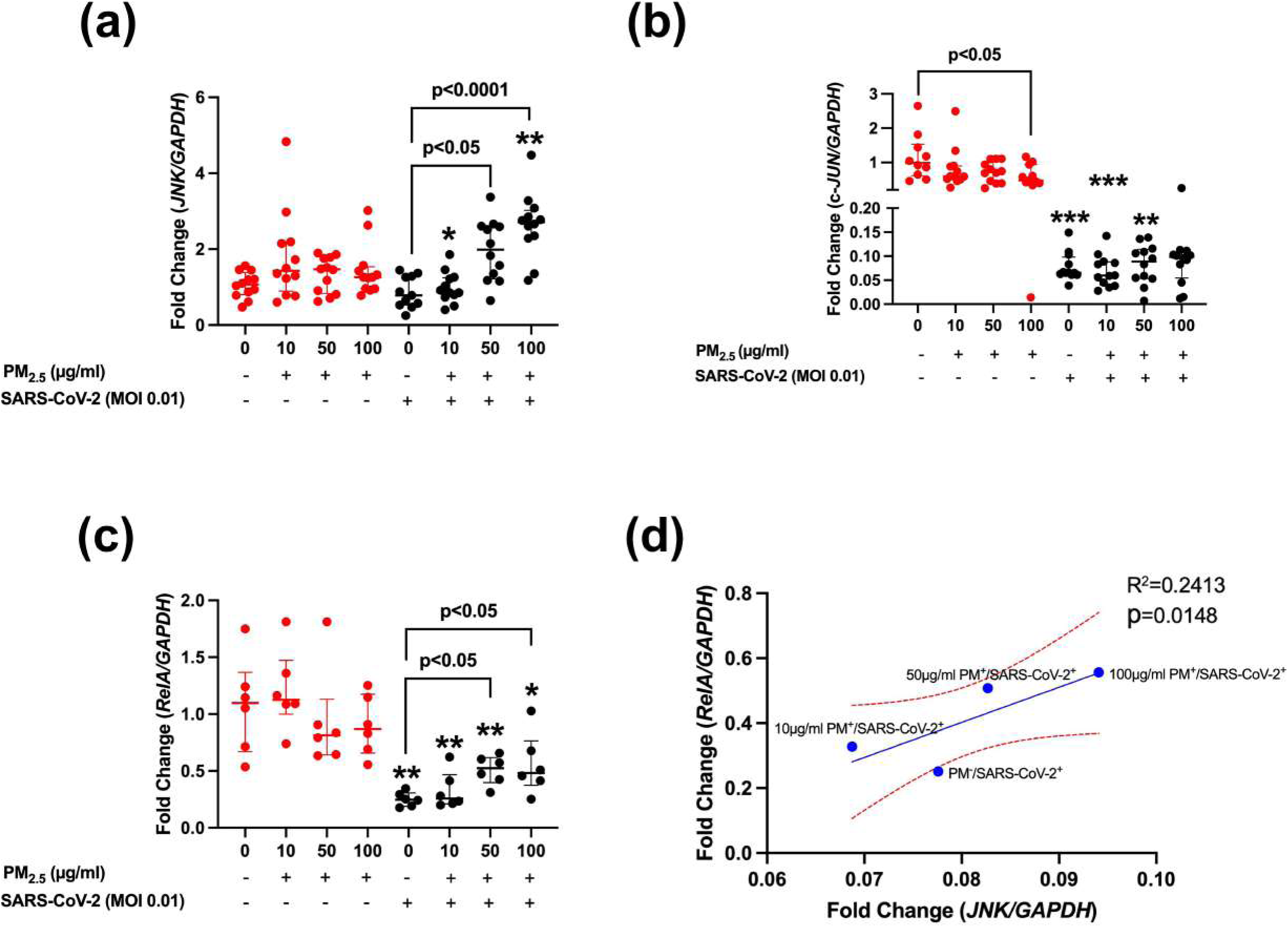
Effects of SARS-CoV-2 and particle matter ≤ 2.5µm (0-100µg/m) on mRNA expression for (a) *MAPK8(JNK)*, (b) *c-JUN* (c) RELA in Calu-3 cells (*p<0.05, **p<0.01, and ***p<0.001 vs corresponding PM_2.5_ concentrations without SARS-CoV-2). (d) The correlation between mRNA expressions of *NF-*κ*B p65 (*RELA) and *JNK*.

Our results demonstrated that although RELA mRNA expression was not altered by PM_2.5_ concentrations used (10-100 µg/ml), in the presence of SARS-CoV-2, *RELA* expression was increased by 50 and 100 µg/ml PM_2.5_ (p<0.05) compared to SARS-CoV-2 alone. On the other hand, SARS-CoV-2 led to a significant decline in the expression of *RELA* in both the absence and presence of 10, 50, or 100 µg/ml PM_2.5_, as compared to the SARS-CoV-2 negative corresponding cultures (**Figure 5c**).

Gene interaction analysis (Genemania.org) has revealed the relation between *RELA*, *JUN, and MAPK8 (JNK)* (**Supplementary** Figure 1) and correlation analysis demonstrating that the mRNA expression for *RELA* and *JNK* was positively correlated by PM_2.5_ and SARS-CoV-2 (R^2^=0.2413, p=0.015) (**Figure 5d**).

### 3.6. Effects **of** PM_2.5_ and SARS-CoV-2 on the mRNA Expression of inflammasome-related genes *NLRP3* and *CASP1* in Calu-3 cells

The expression of NLR family pyrin domain-containing 3 (*NLRP3)* and *Caspase-1 (CASP1)* genes related to the inflammasome was studied to examine the impact of PM_2.5_ and SARS-CoV-2 on the inflammasome response of Calu-3 cells. PM_2.5_ did not affect mRNA expression for these genes; however, 10µg/ml PM_2.5_ significantly reduced the mRNA expression for *NLRP3* in the presence of SARS-CoV-2. Similarly, 50 (p<0.05) and 100 (p<0.001) µg/ml PM_2.5_ reduced CASP1 mRNA expression in the presence of SARS-CoV-2. When the cells treated with PM_2.5_ alone and PM_2.5_ + SARS-CoV-2 were compared, we found that *NLRP3* mRNA expression induced by 0 (p<0.001), 10 (p<0.05), 50 (p<0.05) and 100 (p<0.001) µg/ml PM_2.5_ + SARS-CoV-2 was significantly lower compared to corresponding concentrations of PM_2.5_ alone (**Figure 6a**). In contrast, mRNA expression for CASP1 was significantly enhanced by 0 (p<0.01) and 10 (p<0.05) µg/ml PM_2.5_, as compared to PM_2.5_ alone (0 and 10µg/ml, respectively) (**Figure 6b**).

**Figure 6.**
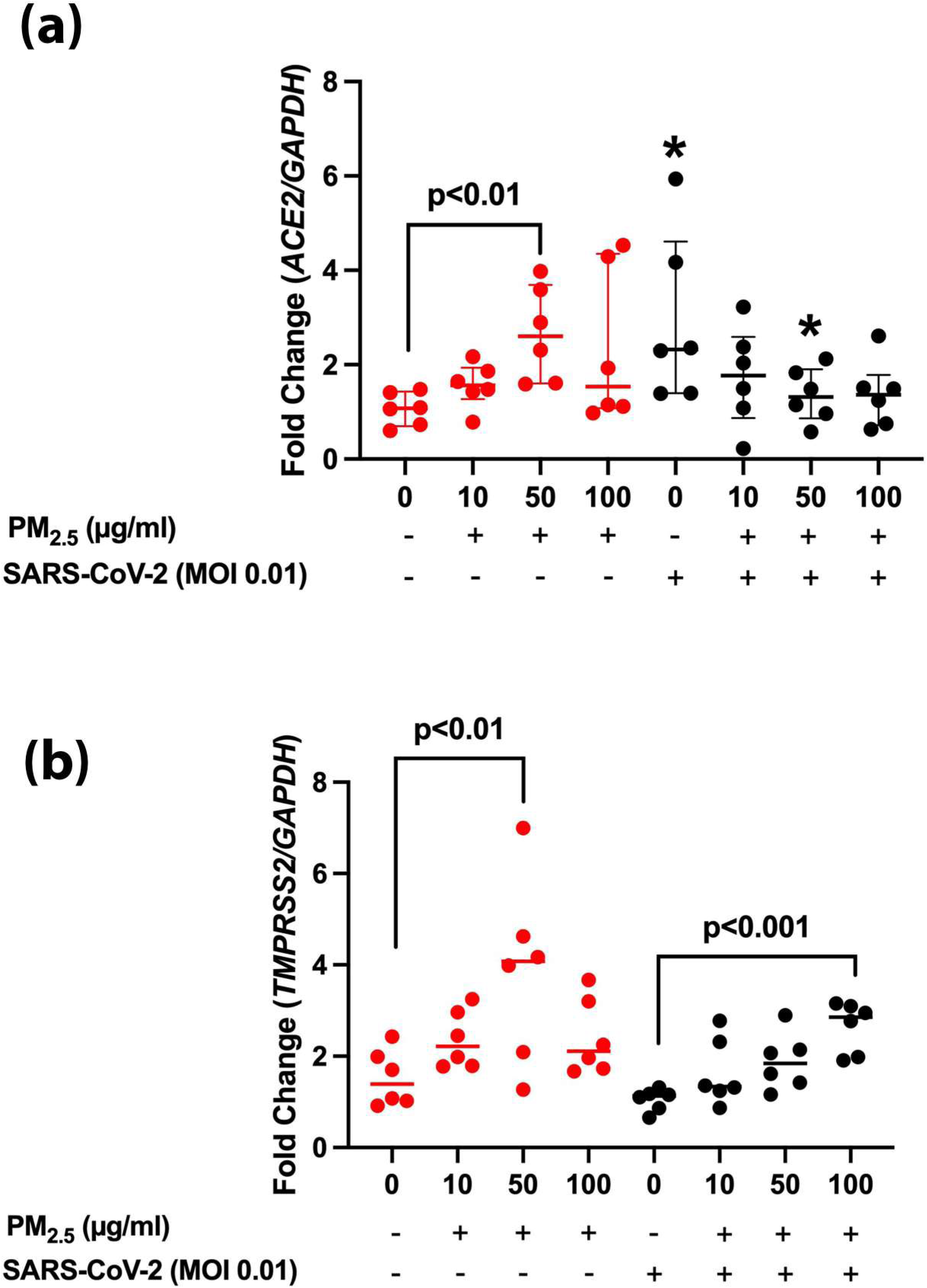
The effect of PM particulate matter ≤ 2.5µm (PM_2.5_) and SARS-CoV-2 on mRNA expression for (a) *NLRP3*, (b) *CASP-1 and (d) IL-1*β in Calu-3 cells (*p<0.05, **p<0.01 vs corresponding PM_2.5_ concentrations); (c) the correlation between mRNA expression for *Casp-1* and *NF-*κ*B p65 (*RELA) following incubation with 10-100 µg/ml PM_2.5_ + SARS-CoV-2.

The correlation analysis showed that the mRNA expression for CASP1 and RELA was correlated by PM_2.5_ and SARS-CoV-2 (R^2^=0.5476, p<0.0001), and this analysis showed that although *RELA* expression was increased, the expression of *CASP-1* was decreased (**Figure 6c**).

The mRNA levels of IL-1β were significantly increased in 10 (p<0.001) and 100 ug/ml PM_2.5_ (p<0.01) exposed Calu-3 cells in absence of SARS-CoV-2. Expression of this gene was increased in the cells exposed to SARS-CoV-2 alone compared to the corresponding virus-free group (p<0.01). In the presence of SARS-CoV-2, 100 µg/ml PM_2.5_ significantly decreased *IL-1*β mRNA expression compared to the virus-only group. The mRNA expression was significantly higher in 10 and 50 µg/ml PM_2.5_-exposed groups in the presence of SARS-CoV-2 compared to the corresponding group without virus exposure (**Figure 6d**).

### 3.8. Effects **of** PM_2.5_ and SARS-CoV-2 on IL-6 and GM-CSF mRNA and protein expression in Calu-3 cells

The release of IL-6 was increased in 50 (p<0.001) and 100 (p<0.01) µg/ml of PM_2.5_ groups. Similarly, SARS-CoV-2 significantly induced the release of IL-6 in the presence of 10 µg/ml PM_2.5_ (p<0.0001), as well as co-incubation of PM2.5 and SARS-CoV-2 increased release of IL-6 in all PM_2.5_ concentration compared to the corresponding concentrations of PM_2.5_ alone (p<0.01) (**Figure 7a**). In the absence of SARS-CoV-2, 50 µg/ml and 100 µg/ml PM_2.5_ significantly increased IL6 mRNA levels. Expression of this gene was also significantly increased in Calu-3 cells exposed to SARS-CoV-2 alone compared to the corresponding virus-free group. In the presence of SARS-CoV-2, exposure to 100 µg/ml PM_2.5_ significantly decreased IL-6 mRNA expression compared to the virus-only group. IL-6 mRNA expression was significantly higher in PM_2.5_-exposed groups in presence SARS-CoV-2 compared to their corresponding groups without virus exposure (**Figure 7b**).

**Figure 7.**
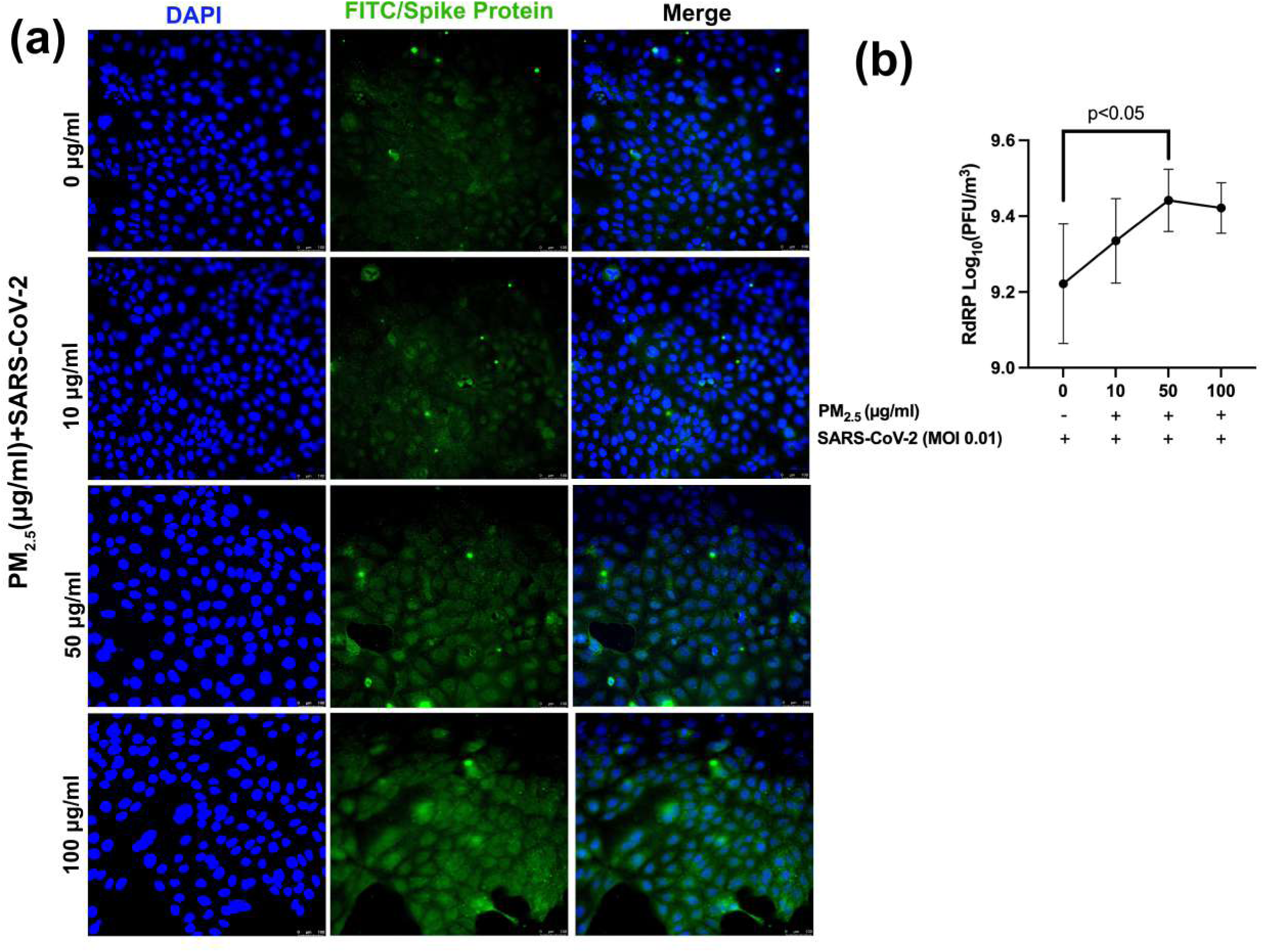
The effects of particulate matter ≤ 2.5µm (PM_2.5_) and SARS-CoV-2 on the protein and mRNA levels of (a and b) and (c and d) GM-CSF from Calu-3 cells following 24 hours’ incubation (**p<0.01, ***p<0.001 vs. corresponding PM_2.5_ concentrations).

On the other hand,the GM-CSF level was elevated by 10 and 100 µg/ml of PM_2.5_, as compared to 0 µg/ml PM_2.5_ (p<0.05). Similarly, 50 µg/ml PM_2.5_ + SARS-CoV-2 increased the release of this cytokine compared to SARS-CoV-2 alone (p<0.01), as well as to the same concentration of 50 µg/ml PM_2.5_ alone (p<0.01), also only SARS-CoV-2 led to increasing GM-CSF level compared to 0 µg/ml PM_2.5_ (p<0.05) (**Figure 7c**). In the absence of SARS-CoV-2, 100 µg/ml PM_2.5_ levels significantly decreased GM-CSF mRNA levels in Calu-3 cells. Expression of this gene did not change in the cells exposed to SARS-CoV-2 alone compared to the corresponding virus-free group. In the presence of SARS-CoV-2, 50 and 100 µg/ml PM_2.5_ concentrations significantly decreased GM-CSF mRNA expression compared to the virus-only group. The mRNA expression was significantly higher in 50ug/ml PM_2.5_-exposed group in presence of SARS-CoV-2 compared to the corresponding group without virus exposure (**Figure 7d**).

### 3.9. Effects of PM_2.5_ and SARS-CoV-2 on IL-8 mRNA and protein expression in Calu-3 cells

Although PM_2.5_ itself did not cause a change in IL-8 release, 10 (p<0.01) and 50 (p<0.001) µg/ml PM_2.5_ decreased the release of this cytokine in the presence of SARS-CoV-2. On the other hand, SARS-CoV-2 itself led to a significant increase in the release of IL-8 (p<0.01), and the release by SARS-CoV-2 + 100µg/ml PM_2.5_ was higher than the same dose of PM_2.5_ alone (p<0.05) (**Figure 8a)**.

**Figure 8.**
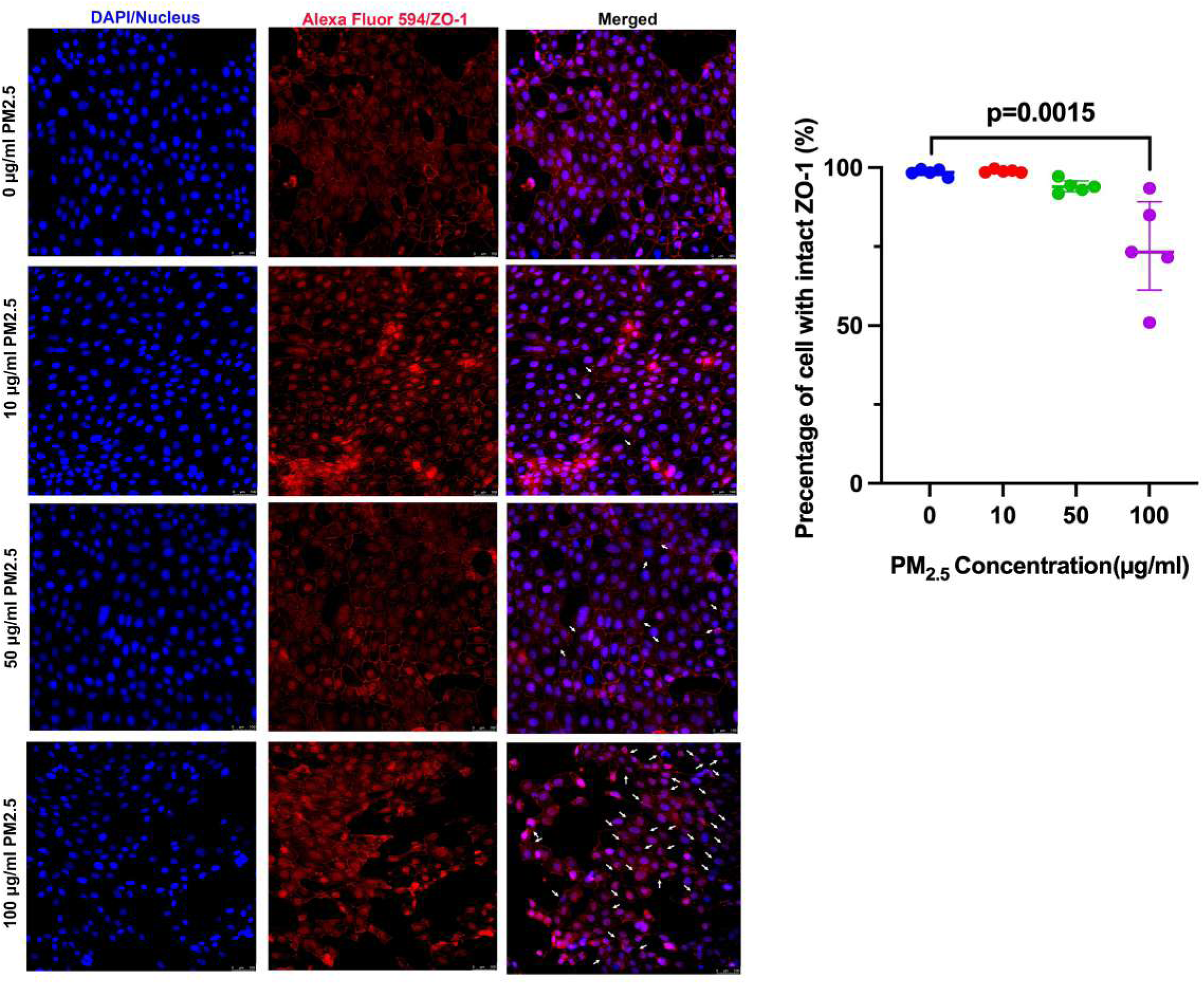
The effects of particulate matter ≤ 2.5µm (PM_2.5_) and SARAS-CoV-2 on release of (a) IL-8, and (b) mRNA expression for IL-8(CXCL-8) (*p<0.05, **p<0.01 vs corresponding PM_2.5_ concentrations); (c) the correlation between mRNA expression for *Casp-1* and *CXCL-8* following incubation with 10-100 µg/ml PM_2.5_ + SARS-CoV-2.

Treating Calu-3 cells with PM_2.5_ did not change the mRNA expression of CXCL8; however, the expression of this gene was significantly increased by SARS-CoV-2 alone (p<0.01). Similarly, SARS-CoV-2 and PM_2.5_ (10-100µg/ml) together led to a significant increase in CXCL8 mRNA expression comparing to corresponding concentrations of PM_2.5_ alone (p<0.01) (**Figure 8b**).

When the correlation analysis was performed, the mRNA expression for CXCL8 and CASP-1 was correlated by PM_2.5_ and SARS-CoV-2 (R^2^=0.3197, p=0.004), and the expression of both genes was reduced by PM_2.5_ (**Figure 8c**).

### 3.10. Effects of PM 2.5 and SARS-CoV-2 on IFNG mRNA expression in Calu-3 cells

IFNG mRNA expression was significantly reduced in Calu-3 cells stimulated with 100 µg/ml PM2.5 (p<0.01). No significant effect was observed at other PM_2.5_ concentrations. Expression of this gene was inhibited by SARS-CoV-2 alone. IFNG expression in Calu-3 cells was completely inhibited in the presence of SARS-CoV-2 at all doses of PM_2.5_. No change was observed with increasing PM_2.5_ concentration. (**Figure 9**)

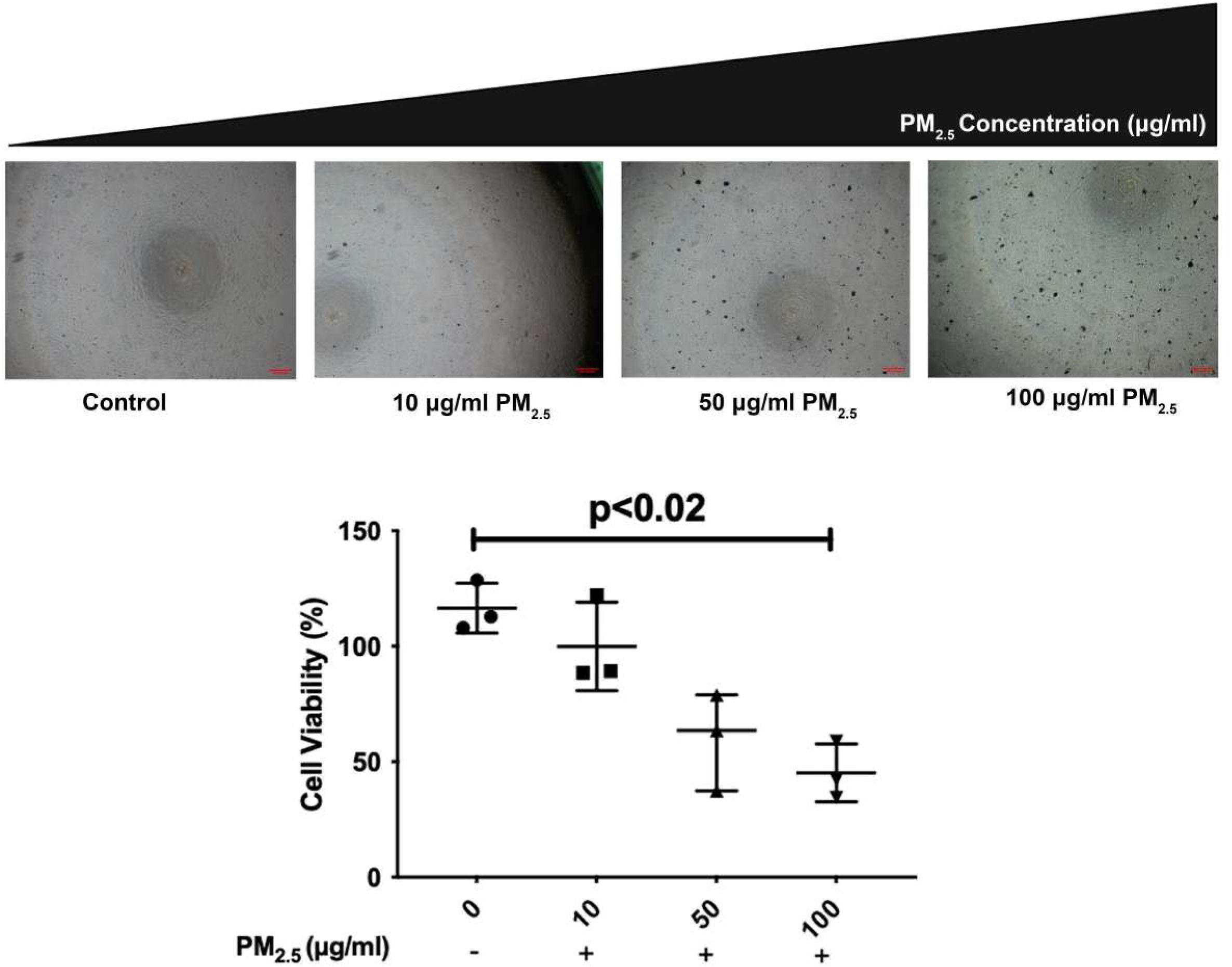

## 4. Discussion

In the current study, we investigated the effects of PM_2.5_ exposure SARS-CoV-2 infection in Calu-3 cells, focusing cellular viability, epithelial barrier integrity, viral entry, antiviral and inflammatory responses. Our findings demonstrated that PM_2.5_, particularly at higher concentrations (100 µg/ml), significantly reduced cell viability, impaired epithelial integrity, and facilitated viral entry by increasing *ACE2* and *TMPRSS2 expression.* Furthermore, both PM_2.5_ and SARS-CoV-2 altered the inflammatory response of Calu-3 cells, which was associated with changes in the inflammasome and antiviral response. These findings suggest that PM_2.5_ can induce the entry of SARS-CoV-2 into airway epithelial cells. PM_2.5_ also, in combination with SARS-CoV-2, can result in decreased antiviral response in host cells, suppressing inflammasome-associated gene expressions, and stimulating the inflammatory response. PM_2.5_ has been shown to exert its detrimental effects on airways by deteriorating respiratory epithelial integrity and cell viability [24, 25]. PM_2.5_ treatment of BEAS-2B epithelial cells led to tight junction disruption by attenuating junction proteins such as Zonula occludens (ZO)-1 and E-cadherin [26]. These findings are in accordance with our findings demonstrating that higher concentration of PM_2.5_ decreased cell viability and induced the disruption in ZO-1 junction of Calu-3 cell cultures. Interestingly, the decrease in ZO-1 expression was associated with increased expression of RdRP gene indicating an increased cellular entry of SARS-CoV-2 in Calu-3 epithelial cells.

Studies have suggested that PM_2.5_ can induce SARS-CoV-2 infection by upregulating the epithelial cell upregulation of ACE2 and TMPRSS2, which are the main receptors mediating SARS-CoV-2 entry to the host cells [27, 28]. Indeed, PM_2.5_ treatment of murine lung enhanced the expression of ACE2 and TMPRSS2 in alveolar type 2 cells (29) Sagawa, T et al, 2021).

Our study showed that 50µg/ml PM_2.5_ induced the mRNA expression for *ACE2* and *TMPRSS2* in Calu-3 cells. Although the co-incubation of a higher concentration of 100µg/ml PM_2.5_ and SARS-CoV-2 enhanced expression of *TMPRSS2,* this did not alter the expression of ACE2, as compared to SARS-CoV-2 alone. However, in immunofluorescence studies, the expression of SARS-CoV-2 S protein showed an increasing trend following co-incubation of Calu-3 cells with SARS-CoV-2 and higher concentrations of 50-100µg/ml PM_2.5_ that was associated with RdRP mRNA expression. Together, these findings suggest an increased viral replication by PM_2.5_ in Calu-3 cells.

Viral spike (S) protein, which contains two subunits (S1 and S2), is associated with viral entry by binding to the host cell’s ACE2 receptor, and after entering the cells, TMPRSS2 cleaves the S2 subunit to activate and mature the virus [29]. Non-structural proteins (NSPs) also have a crucial effect on interrupting host replication mechanisms and regulating the assembly of newly synthesized viral products. They also adversely affect the innate immune response against the virus [30]. There are 16 NSPs in the SARS-CoV-2 genome with different roles; the most important NSPs are NSP12, also known as RdRP, and nsp13 (helicase), which are involved in directing viral genomes and protein synthesis [31].

On the other hand, although SARS-CoV-2 alone increased ACE2 mRNA expression in Calu-3 cells, the co-incubation of Calu-3 cells with both 50µg/ml PM_2.5_ and SARS-CoV-2 led to a decrease in *ACE2* mRNA expression comparing to PM2.5 alone. Similarly, Gao et al. showed that ACE2 expression could be downregulated in SARS-CoV-2 infected cells by the virus’s S protein, which in turn could disrupt the renin-angiotensin system (RAS) homeostasis, leading to an increase in angiotensin II (AngII) and pulmonary vascular damage in patients with COVID-19 [32].

Interestingly, it has been suggested that PM_2.5_-induced ACE2 expression could lead to increased viral load and, subsequently, depletion of ACE2. The occupation of ACE2 by SARS-CoV-2 binding can cause downregulation of the receptor that may lead to the induction of AngII axis resulting in an inflammatory response [12]. ACE2 is reported to be responsible for maintaining ACE/ACE2 and Ang-II/angiotensin-(1–7) balance and has an anti-inflammatory effect; thus, in addition to disrupting RAS hemostasis, the downregulation of ACE2 by SARS-CoV-2 may lead to increasing inflammation and organ damage [33].

The innate immune system is the major defense mechanism against viral invasion. It works through the interaction of pattern recognition receptors (PRRs) and pathogen-associated molecular patterns (PAMPs) to activate inflammatory responses and programmed cell death to suppress viral infection and replication [19]. The excessive response of the innate immune system can cause serious inflammatory reactions and tissue damage known as cytokine storm, a commonly agreed severe COVID-19 characteristic [34, 35]. Cytokine storm is an excessive and recurrent inflammatory response mainly caused by immune cells. Major cytokines in this exacerbated inflammatory response are IL-1β, IL-6, IL-8, colony-stimulating factors (CSF), and granulocyte macrophage (GM)-CSF [34]. Tapas Patra and co-workers showed that S protein of SARS-CoV-2 induced IL-6 release by regulating NF-κB and AP-1 pathways [36]. GM-CSF was also upregulated in severe COVID-19 conditions such as acute respiratory distress syndrome (ARDS) [37].

We demonstrated that PM_2.5_ led to increased IL-6 and GM-CSF release from Calu-3 cells both in the absence and presence of SARS-CoV-2. However, although SARS-CoV-2 enhanced the release of IL-8, PM_2.5_ alone did not show an effect, and when co-incubated with SARS-CoV-2, these particles decreased both mRNA and protein expression of IL-8. Few studies have focused on the mechanistic effect of PMs on cytokine production in SARS-CoV-2 infection. Damariz Marín-Palma et al. reported that PM_10_ could induce the production of IL-6 in PBMCs from healthy individuals incubated with SARS-CoV-2, which was associated with increased viral replication [38]. Also Zhang et al confirmed that SARS-CoV-2 S protein receptor binding domain induce the IL-6 and IL-8 release in human bronchial epithelial cells [39]. In another study, it was shown that 24 hours of heat-inactivated SARS-CoV-2 (2.5 × 10^2^ genome copies/cm^2^) co-exposure increased IL-6 and IL-8 release after 72 hours of pre-exposure with low concentration PM_2.5_ (2.5 µg/cm^2^) in A549 cells. In the same study, it was observed that IL-6 release was increased in all exposure groups after 72 hours of exposure of only PM_2.5_, SARS-CoV-2 alone, and PM_2.5_ and SARS-CoV-2 together, compared to the untreated control group, while IL-8 release was increased by PM_2.5_ and SARS-CoV-2 co-exposure it has only been shown to increase with respect to the group exposed to PM_2.5_ [40]. In a recent study, when SARS-CoV-2 pseudovirus and PM_2.5_ were exposed to ACE2 transgenic mice, it was reported that IL-6 and TNFα cytokine release increased in BAL fluids compared to the pseudovirus-only group that did not receive PM_2.5_ [41].

While excessive activation of the innate immune system and overproducing IL-8 can threaten the patient’s health, any disruption in the innate immune system can cause a deficiency against microbial and viral infection Chang et al found that SARS-CoV S protein-induced IL-8 promoter activity was inhibited by the specific inhibitors of MAPK cascades and they suggested that S protein of SARS-CoV could induce release of IL-8 in the lung cells via activations of MAPKs and AP-1[42]. Thus, any disruption in this process can cause increased viral replication and infection in host cells [43]. For the first time, our results showed that IL-8 protein and mRNA expression, which was increased in only the SARS-CoV-2 incubated group, gradually declined following an increase in PM_2.5_ concentration in PM_2.5_ and SARS-CoV-2 co-incubated groups. To understand the underlying mechanisms, at the first step, we identified the most related genes in regulating *CXCL-8* expression using the Genemania database (**Supplementary** Figure 2), and according to bioinformatic analysis, the most associated genes that correlated with *CXCL-8* were *c-JUN/MAPK8 (JNK)* and *NF-*κ*B p65 (RelA)*.

NF-κB composed of different subunits such as NF-κB1 p50, NF-κB2 p52, RELA p65, RelB, and c-Rel, which is critical in governing inflammatory responses by activating various pro-inflammatory genes and cytokines [44]. JNK, on the other hand, can regulate the activity of immune cells and the induction of pro-inflammatory cytokines like IL-1β, IL-1α, IL-2, IL-6, IL-8, and TNF-α [45, 46]. In cells that overexpress the SARS-CoV spike protein, JNK phosphorylation is facilitated by protein kinase C epsilon, and IL-8 production is contingent upon AP-1 activity [47]. Both JNK and NF-κB have a crucial role in the inflammatory response against SARS-CoV-2, and our results also showed an increase in mRNA expression of *JNK* and *NF-*κ*B p65 (*RELA) in the presence of both PM_2.5_ and SARS-CoV-2. In line with our findings, Su et al. reported that SARS-CoV-2 Open Reading Frame (ORF) 7a induced NF-κB activation and expression of inflammatory cytokines [48]. Furthermore, Forsyth and his co-workers confirmed that SARS-CoV-2 S1 protein could activate JNK and NF-κB in human lung epithelial cells [49]. JNK phosphorylation was also observed in cells infected with MHV or SARS-CoV and in cells overexpressing the N, 3a, 3b, or 7a proteins of SARS-CoV. Also, JNK and Akt are essential for the establishment of persistent SARS-CoV infection[47] . C-Jun expression, an essential element of the JNK/AP pathway-1 pathway plays a crucial role in the IRE-1α-mediated transcriptional control of stress response genes that possess anti-inflammatory and cytoprotective functions [50]. Rasmussen and his co-workers have reported that *JNK* and *NF-*κ*B p65* can inversely affect the expression of IL-8 [51].

In a study by Marcetti et al. (2024), it was reported that both 72 hours of combined exposure to PM and SARS-CoV-2, and 24 hours of exposure to SARS-CoV-2 after 72 hours of PM exposure, resulted in decreased NF-κB protein levels only in SARS-CoV-2, while PM exposure increased expression. The decrease in NF-κB protein levels in A549 cells exposed only to the virus in this study is consistent with the decrease in RELA observed in our experiments. However, in our study, RELA gene expression was reduced in the presence of virus at all concentrations of PM_2.5_. Marcetti et al. reported that NF-κB protein levels increased at a concentration of 2.5 µg/cm^2^ PM2.5 (approximately 12 µg/ml per well for a 6-well plate).

According to the confirmed role of *NF-*κ*B p65/* RELA in controlling *CXCL-8* and inflammasomal response (**Supplementary** Figure 3) [52–54], we presumed an inflammasomal response as a linker between *NF-*κ*B* and *CXCL-8* expression. Inflammasomes are protective immune responses that involve damage responses such as microbial and viral infection through the innate immune system [55]. The most important parts of the inflammasome complex are NLRPs (NLRP1, NLRP3) and cysteine proteases called CASP-1, which govern inflammation or cell death by activating inflammatory cytokines like IL-1β, IL-18, and IL-8 [55–57]. The inflammasome is an important part of the inflammatory response against SARS-CoV-2. Rodrigues et al. have confirmed the role of the inflammasome in COVID-19 pathophysiology and the correlation between inflammasomal products like *CASP-1* and COVID-19 severity [58]. However, the inflammasomal response modifications are not clearly understood in the presence of SARS-CoV-2 and PM_2.5_. Our results showed that, while only PM_2.5_ did not alter the *NLRP3* and *CASP-1*, co-incubation of PM_2.5_ with SARS-CoV-2 decreased NLRP3 expression, especially in the presence of 10µg/ml PM_2.5_. On the other hand, *CASP-1*, which was overexpressed after only SARS-CoV-2 incubation, started to decrease by increased concentrations of PM_2.5._

Besides the inflammatory regulatory role of *NF-*κ*B*, it is a major signaling pathway that can transcriptionally regulate inflammasomes. However, this interaction is complex, while studies have shown the importance of *NF-*κ*B* activation of the priming step for inflammasomal response. Zhong et al. in contrast, reported that the activation of *NF-*κ*B* led to inhibition of inflammasomal response through p62 activation [44, 59]. Also, our gene expression and correlation analysis for *CASP-1* and RELA indicated that along with increasing PM_2.5_ concentrations and upregulation of RELA in the PM_2.5_ and SARS-CoV-2 incubated groups, *CASP-1* mRNA expression declined and correlated with RELA **(Figure 6c)**.

As previously described, our results have shown a slight decrease in IL-8 protein and mRNA expression in PM_2.5_ and SARS-CoV-2 incubated groups. The regulatory role of *CASP-1* on IL-8 expression has been confirmed previously, Koenen et al. have stated that inhibition of intrinsic *CASP-1* activity can cause diminished IL-8 production, also using pan-caspase inhibitor, z-VAD-fmk results in a reduction of TNFα, IL-6, and IL-8 expression [60, 61]. Also, Mortaz et al. have confirmed that inhibition of CASP-1 results in downregulating cigarette smoke-induced CXCL-8 expression in human bronchial epithelial cells (HBE-14o) [57]. To confirm the relation between IL-8 production and mRNA expression with inflammasomal response in our study, we also performed gene expression and correlation analysis for *CASP-1* and *CXCL-8*, according to our results, along with increasing PM_2.5_ concentration in SARS-CoV-2 incubated groups, both *CXCL-8* and *CASP-1* mRNA expression attenuated, and this decreasing was significantly correlated (**Figure 8c**).

IFNG (Interferon gamma), a key component of the antiviral response, is reduced by both PM2.5 alone (100 μg/ml, p<0.01) and SARS-CoV-2 (p<0.01), and its expression is completely suppressed by their combined exposure. This synergistic effect clearly demonstrates that PM2.5’s inhibition of IFNG expression contributes to the suppression of the antiviral response.

Overall, our findings suggest that PM2.5 exposure enhances viral entry and replication (ACE2/TMPRSS2 regulation, Spike/RdRP upregulation), alters antiviral defense mechanisms (IFNG), and suppresses some inflammatory pathways (CASP-1). This unbalanced effect provides an explanation for the potential role of air pollution in increasing the spread of SARS-CoV-2 infection.

Limitations of our study include the lack of an accurate assessment of inflammasome performance by measuring IL-1β, IL-18, and granzyme D levels. Furthermore, we rely on the transcription and phosphorylation of JNK, JUN, and NFKB, which are important for inflammatory signaling pathways. Future studies should include longer incubation times and direct measurement of this cytokine/protein expression.

## 5. Conclusion

In conclusion, our results demonstrated that PM_2.5_ led to decreased cell viability, disrupted epithelial cells barrier integrity, and increased viral entry via upregulating *ACE2* and *TMPRSS2* and could modulate the innate immune response in SARS-CoV-2 infected Calu-3 cells. These findings suggest a possible role of PM_2.5_ in modulating molecular mechanisms like inflammasomal response involved in innate immune response disruption against SARS-CoV-2 infection.

## Author Contributions

Conceptualization, O.K. and HB.; methodology, O.K., H.R., G.E., F.C., and HB.; software, O.K. and H.R.; validation, O.K., H.R., and H.B.; formal analysis, O.K., H.R., G.E., F.C., and H.B.; investigation, O.K., H.R., G.E., F.C., and H.B.; resources, O.K., H.R., G.E., F.C., and H.B.; data curation, O.K., H.R. writing—original draft preparation, O.K. and H.R.; writing—review and editing, O.K., H.R., and H.B.; visualization, O.K., H.R.; supervision O.K., and H.B.; project administration, O.K. and H.B.; funding acquisition, O.K. All authors have read and agreed to the published version of the manuscript.

## Data Availability Statement

Data are contained within the article. Data is available on request from the authors.

## Acknowledgments

, The authors acknowledge the use of the services and facilities of the Koç University Research Center for Translational Medicine (KUTTAM), funded by the Presidency of Turkey, Head of Strategy and Budget.

## Legends for supplementary figures

**Figure S1:** Illustration showing the interaction of NF-κB p65 (RELA) with MAPK 8(JNK) and c-JUN.

**Figure S2:** Results from Genemania.org database have shown the interaction among IL-8 with the RELA, c-JUN, and MAPK8.

**Figure S3:** Data from the Genemania.org database has shown the interaction of CASP1 with RELA and *IL-8*.

**Supplementary Table 1.**
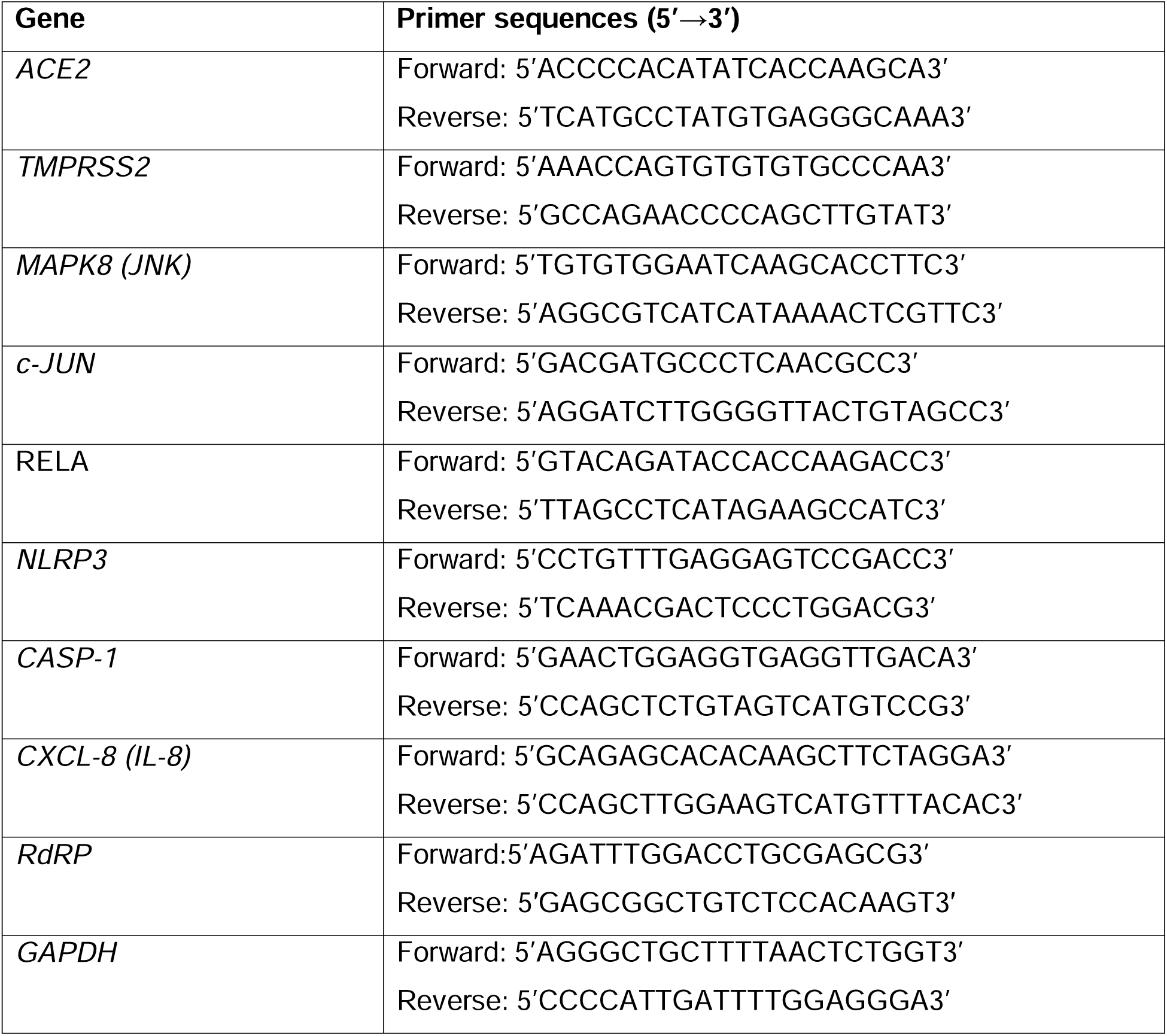
Primers used for real-time PCR.

## References

1. Gómez, C.E., B. Perdiguero, and M. Esteban, Emerging SARS-CoV-2 variants and impact in global vaccination programs against SARS-CoV-2/COVID-19. Vaccines, 2021. 9(3): p. 243.

2. Rajabi, H., et al., Forthcoming complications in recovered COVID-19 patients with COPD and asthma; possible therapeutic opportunities. Cell Communication and Signaling, 2022. 20(1): p. 173.

3. Mandal, C.C., et al., Combinatorial influence of environmental temperature, obesity and cholesterol on SARS-CoV-2 infectivity. Scientific Reports, 2022. 12(1): p. 4796.

4. Michalakis, K. and I. Ilias, SARS-CoV-2 infection and obesity: Common inflammatory and metabolic aspects. Diabetes & Metabolic Syndrome: Clinical Research & Reviews, 2020. 14(4): p. 469–471.

5. Gong, Z., et al., Natural and socio-environmental factors in the transmission of COVID-19: a comprehensive analysis of epidemiology and mechanisms. BMC Public Health, 2024. 24(1): p. 2196.

6. Meyerowitz, E.A., et al., Transmission of SARS-CoV-2: a review of viral, host, and environmental factors. Annals of internal medicine, 2021. 174(1): p. 69–79.

7. Brown, K.L., et al., Estimating the heritability of SARS-CoV-2 susceptibility and COVID-19 severity. Nature Communications, 2024. 15(1): p. 367.

8. Lembo, R., et al., Air pollutants and SARS-CoV-2 in 33 European countries. Acta Bio Medica: Atenei Parmensis, 2021. 92(1).

9. Wu, X., et al., Air pollution and COVID-19 mortality in the United States: Strengths and limitations of an ecological regression analysis. Science advances, 2020. 6(45): p. eabd4049.

10. Semczuk-Kaczmarek, K., et al., Association between air pollution and COVID-19 mortality and morbidity. Internal and emergency medicine, 2022: p. 1–7.

11. Zhang, J., et al., Long-term exposure to air pollution and risk of SARS-CoV-2 infection and COVID-19 hospitalisation or death: Danish nationwide cohort study. European Respiratory Journal, 2023. 62(1).

12. Bayram, H., et al., Issue 4—Impact of air pollution on COVID-19 mortality and morbidity: An epidemiological and mechanistic review. Pulmonology, 2024.

13. Hoffmann, M., et al., SARS-CoV-2 cell entry depends on ACE2 and TMPRSS2 and is blocked by a clinically proven protease inhibitor. cell, 2020. 181(2): p. 271–280. e8.

14. Riva, D., et al., Low dose of fine particulate matter (PM2. 5) can induce acute oxidative stress, inflammation and pulmonary impairment in healthy mice. Inhalation toxicology, 2011. 23(5): p. 257–267.

15. Botto, L., et al., Effects of PM2. 5 exposure on the ACE/ACE2 pathway: possible implication in COVID-19 pandemic. International Journal of Environmental Research and Public Health, 2023. 20(5): p. 4393.

16. Kayalar, Ö., et al., Existence of SARS-CoV-2 RNA on ambient particulate matter samples: a nationwide study in Turkey. Science of the Total Environment, 2021. 789: p. 147976.

17. Domingo, J.L., M. Marquès, and J. Rovira, Influence of airborne transmission of SARS-CoV-2 on COVID-19 pandemic. A review. Environmental research, 2020. 188: p. 109861.

18. Linillos-Pradillo, B., et al., Determination of SARS-CoV-2 RNA in different particulate matter size fractions of outdoor air samples in Madrid during the lockdown. Environmental Research, 2021. 195: p. 110863.

19. Diamond, M.S. and T.-D. Kanneganti, Innate immunity: the first line of defense against SARS-CoV-2. Nature immunology, 2022. 23(2): p. 165–176.

20. Bayram, H., et al., Effect of serum on diesel exhaust particles (DEP)-induced apoptosis of airway epithelial cells in vitro. Toxicology letters, 2013. 218(3): p. 215–223.

21. Schneider, C.A., W.S. Rasband, and K.W. Eliceiri, NIH Image to ImageJ: 25 years of image analysis. Nature methods, 2012. 9(7): p. 671–675.

22. Warde-Farley, D., et al., The GeneMANIA prediction server: biological network integration for gene prioritization and predicting gene function. Nucleic acids research, 2010. 38(suppl_2): p. W214–W220.

23. Livak, K.J. and T.D. Schmittgen, Analysis of relative gene expression data using real-time quantitative PCR and the 2− ΔΔCT method. methods, 2001. 25(4): p. 402–408.

24. Deng, X., et al., PM2. 5-induced oxidative stress triggers autophagy in human lung epithelial A549 cells. Toxicology in vitro, 2013. 27(6): p. 1762–1770.

25. Zhang, H.-h., et al., Physical and chemical characteristics of PM2. 5 and its toxicity to human bronchial cells BEAS-2B in the winter and summer. Journal of Zhejiang University. Science. B, 2018. 19(4): p. 317.

26. Zhao, C., et al., Respiratory exposure to PM2. 5 soluble extract disrupts mucosal barrier function and promotes the development of experimental asthma. Science of The Total Environment, 2020. 730: p. 139145.

27. Wang, B., et al., Is there an association between the level of ambient air pollution and COVID-19? American Journal of Physiology-Lung Cellular and Molecular Physiology, 2020. 319(3): p. L416–L421.

28. Li, H.-H., et al., Upregulation of ACE2 and TMPRSS2 by particulate matter and idiopathic pulmonary fibrosis: a potential role in severe COVID-19. Particle and Fibre Toxicology, 2021. 18: p. 1–13.

29. Jackson, C.B., et al., Mechanisms of SARS-CoV-2 entry into cells. Nature reviews Molecular cell biology, 2022. 23(1): p. 3–20.

30. Tam, D., et al., Targeting SARS-coV-2 non-structural proteins. International Journal of Molecular Sciences, 2023. 24(16): p. 13002.

31. Malone, B., et al., Structures and functions of coronavirus replication–transcription complexes and their relevance for SARS-CoV-2 drug design. Nature Reviews Molecular Cell Biology, 2022. 23(1): p. 21–39.

32. Gao, X., et al., Spike-mediated ACE2 down-regulation was involved in the pathogenesis of SARS-CoV-2 infection. Journal of Infection, 2022. 85(4): p. 418–427.

33. Ni, W., et al., Role of angiotensin-converting enzyme 2 (ACE2) in COVID-19. Critical Care, 2020. 24: p. 1–10.

34. Song, P., et al., Cytokine storm induced by SARS-CoV-2. Clinica chimica acta, 2020. 509: p. 280–287.

35. Codina, H., et al., Elevated anti-SARS-CoV-2 antibodies and IL-6, IL-8, MIP-1β, early predictors of severe COVID-19. Microorganisms, 2021. 9(11): p. 2259.

36. Patra, T., et al., SARS-CoV-2 spike protein promotes IL-6 trans-signaling by activation of angiotensin II receptor signaling in epithelial cells. PLoS pathogens, 2020. 16(12): p. e1009128.

37. Lang, F.M., et al., GM-CSF-based treatments in COVID-19: reconciling opposing therapeutic approaches. Nature Reviews Immunology, 2020. 20(8): p. 507–514.

38. Marín-Palma, D., et al., PM10 promotes an inflammatory cytokine response that may impact SARS-CoV-2 replication in vitro. Frontiers in Immunology, 2023. 14: p. 1161135.

39. Zhang, R.-G., et al., SARS-CoV-2 spike protein receptor binding domain promotes IL-6 and IL-8 release via ATP/P2Y2 and ERK1/2 signaling pathways in human bronchial epithelia. Molecular Immunology, 2024. 167: p. 53–61.

40. Marchetti, S., et al., Shedding light on the cellular mechanisms involved in the combined adverse effects of fine particulate matter and SARS-CoV-2 on human lung cells. Science of the Total Environment, 2024. 952: p. 175979.

41. Lin, M.-W., et al., The influence of PM2. 5 exposure on SARS-CoV-2 infection via modulating the expression of angiotensin converting enzyme II. Journal of Hazardous Materials, 2025. 485: p. 136887.

42. Chang, Y.-J., et al., Induction of IL-8 release in lung cells via activator protein-1 by recombinant baculovirus displaying severe acute respiratory syndrome-coronavirus spike proteins: identification of two functional regions. The Journal of Immunology, 2004. 173(12): p. 7602–7614.

43. Blanco-Melo, D., et al., Imbalanced host response to SARS-CoV-2 drives development of COVID-19. Cell, 2020. 181(5): p. 1036–1045. e9.

44. Liu, T., et al., NF-κB signaling in inflammation. Signal transduction and targeted therapy, 2017. 2(1): p. 1–9.

45. Roy, P.K., et al., Role of the JNK signal transduction pathway in inflammatory bowel disease. World journal of gastroenterology: WJG, 2008. 14(2): p. 200.

46. Chen, N., et al., JNK kinase promotes inflammatory responses by inducing the expression of the inflammatory amplifier TREM1 during influenza a virus infection. Virus Research, 2025: p. 199577.

47. Fung, T.S. and D.X. Liu, Activation of the c-Jun NH2-terminal kinase pathway by coronavirus infectious bronchitis virus promotes apoptosis independently of c-Jun. Cell death & disease, 2017. 8(12): p. 3215.

48. Su, C.-M., L. Wang, and D. Yoo, Activation of NF-κB and induction of proinflammatory cytokine expressions mediated by ORF7a protein of SARS-CoV-2. Scientific reports, 2021. 11(1): p. 13464.

49. Forsyth, C.B., et al., The SARS-CoV-2 S1 spike protein promotes MAPK and NF-kB activation in human lung cells and inflammatory cytokine production in human lung and intestinal epithelial cells. Microorganisms, 2022. 10(10): p. 1996.

50. Bartolini, D., et al., Endoplasmic reticulum stress and NF-kB activation in SARS-CoV-2 infected cells and their response to antiviral therapy. IUBMB life, 2022. 74(1): p. 93–100.

51. Rasmussen, M.K., et al., IL-8 and p53 are inversely regulated through JNK, p38 and NF-κB p65 in HepG2 cells during an inflammatory response. Inflammation Research, 2008. 57: p. 329–339.

52. Yang, Q., et al., RelA/MicroRNA-30a/NLRP3 signal axis is involved in rheumatoid arthritis via regulating NLRP3 inflammasome in macrophages. Cell death & disease, 2021. 12(11): p. 1060.

53. García, J.A., et al., Disruption of the NF-κB/NLRP3 connection by melatonin requires retinoid-related orphan receptor-α and blocks the septic response in mice. The FASEB Journal, 2015. 29(9): p. 3863–3875.

54. Kunsch, C., et al., Synergistic transcriptional activation of the IL-8 gene by NF-kappa B p65 (RelA) and NF-IL-6. Journal of immunology (Baltimore, Md.: 1950), 1994. 153(1): p. 153–164.

55. Guo, H., J.B. Callaway, and J.P. Ting, Inflammasomes: mechanism of action, role in disease, and therapeutics. Nature medicine, 2015. 21(7): p. 677–687.

56. Schroder, K. and J. Tschopp, The inflammasomes. cell, 2010. 140(6): p. 821–832.

57. Mortaz, E., et al., Cigarette smoke induces the release of CXCL-8 from human bronchial epithelial cells via TLRs and induction of the inflammasome. Biochimica et Biophysica Acta (BBA)-Molecular Basis of Disease, 2011. 1812(9): p. 1104–1110.

58. Rodrigues, T.S., et al., Inflammasomes are activated in response to SARS-CoV-2 infection and are associated with COVID-19 severity in patients. Journal of Experimental Medicine, 2021. 218(3).

59. Zhong, Z., et al., NF-κB restricts inflammasome activation via elimination of damaged mitochondria. Cell, 2016. 164(5): p. 896–910.

60. Koenen, T.B., et al., The inflammasome and caspase-1 activation: a new mechanism underlying increased inflammatory activity in human visceral adipose tissue. Endocrinology, 2011. 152(10): p. 3769–3778.

61. Lipinska, K., et al., Applying caspase-1 inhibitors for inflammasome assays in human whole blood. Journal of immunological methods, 2014. 411: p. 66–69.

