## Supplemental Figure 1 for "PM_2.5_ Exposure Facilitates SARS-CoV-2 Infection through ACE2/TMPRSS2 Regulation and Suppression of Anti-Viral Response"

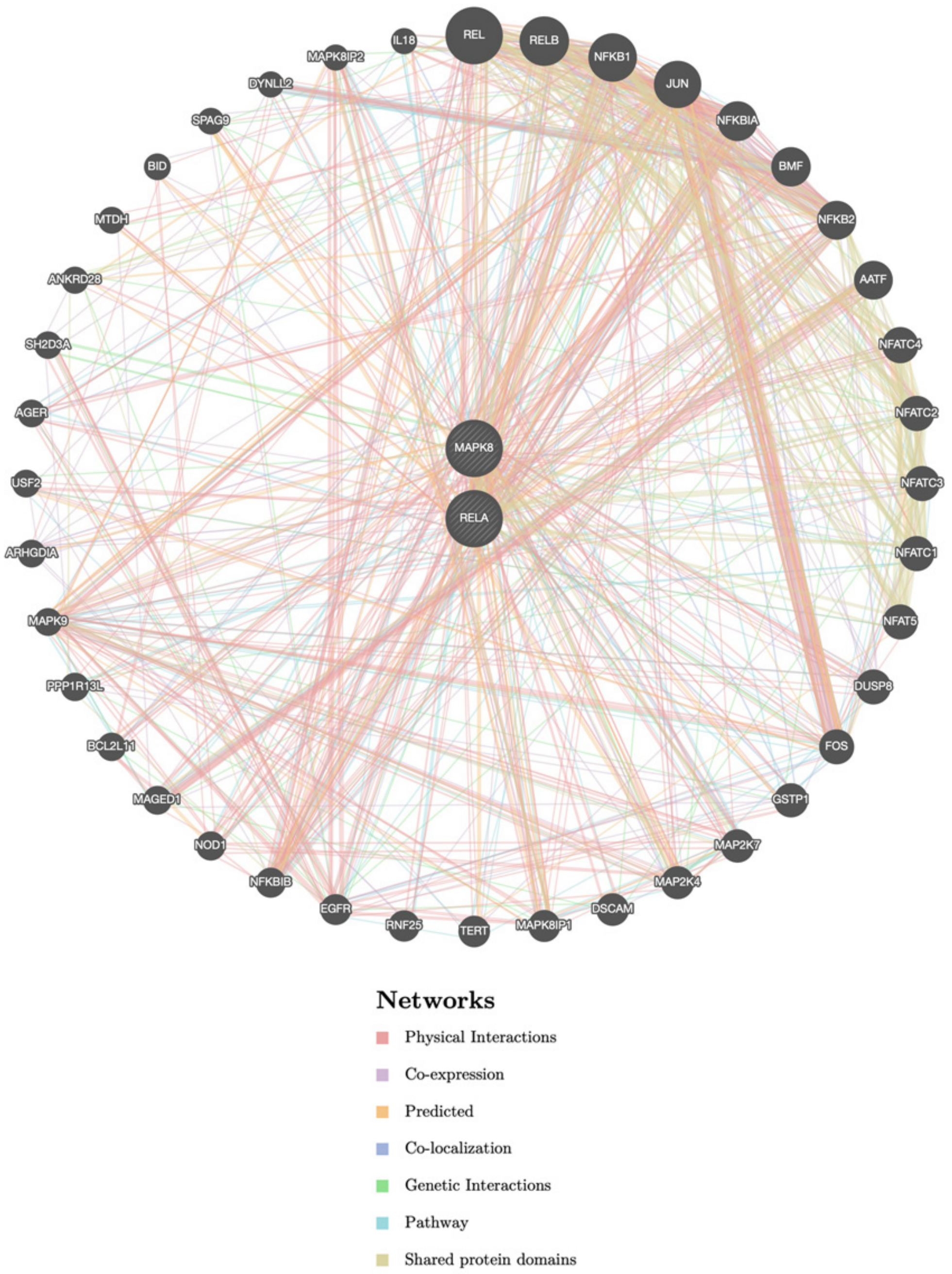


Supplementary figure 1: Illustration showing the interaction between NF-κB p65 (RelA) with MAPK 8(JNK) and c-JUN.

| **RelA** | Score | |
|---|---|---|
| Networks | MAPK8 | c-JUN |
| Physical Interactions | 1 | 0.002 |
| Co-Expression | 0.018 | 0.012 |
| Predicted | 0.02 | 0.015 |
| Co-Localization | - | - |
| Genetic Interactions | 0.002 | 0.25 |
| Pathway | 0.0017 | 0.001 |
| Shared protein domain | - | - |
