## Supplemental Figure 2 for "PM_2.5_ Exposure Facilitates SARS-CoV-2 Infection through ACE2/TMPRSS2 Regulation and Suppression of Anti-Viral Response"

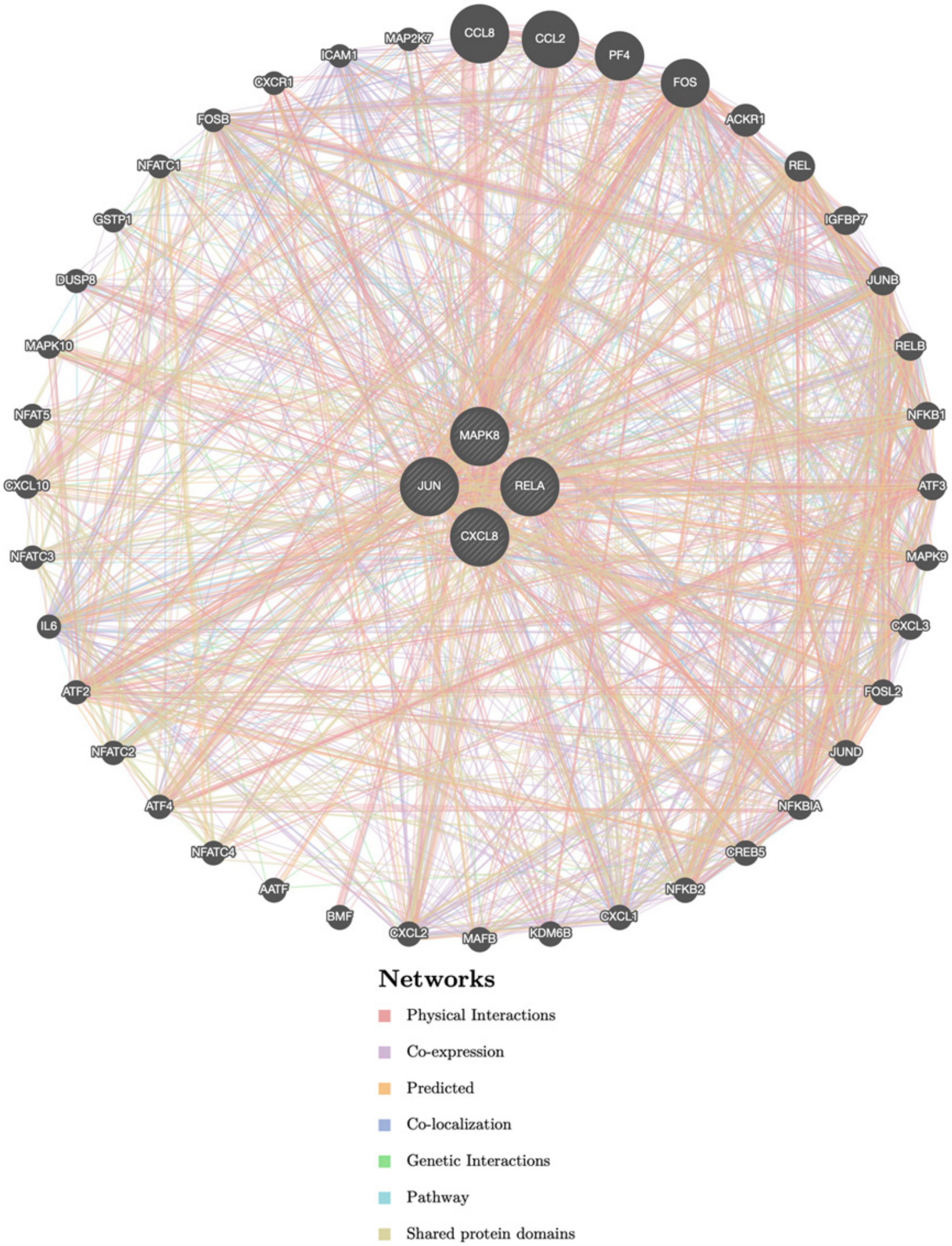


Supplementary Figure 2: Results from Genemania.org database have shown the interaction among CXCL-8 with the RelA, c-JUN, and MAPK8.

| **CXCL-8** | Score | | |
|---|---|---|---|
| Networks | MAPK8 | c-JUN | RelA |
| Physical Interactions | - | - | 0.006 |
| Co-Expression | - | 0.019 | 0.01 |
| Predicted | - | - | - |
| Co-Localization | - | - | 0.01 |
| Genetic Interactions | - | - | - |
| Pathway | - | 0.003 | 0.005 |
| Shared protein domain | - | - | - |
