## Supplemental Figure 3 for "PM_2.5_ Exposure Facilitates SARS-CoV-2 Infection through ACE2/TMPRSS2 Regulation and Suppression of Anti-Viral Response"

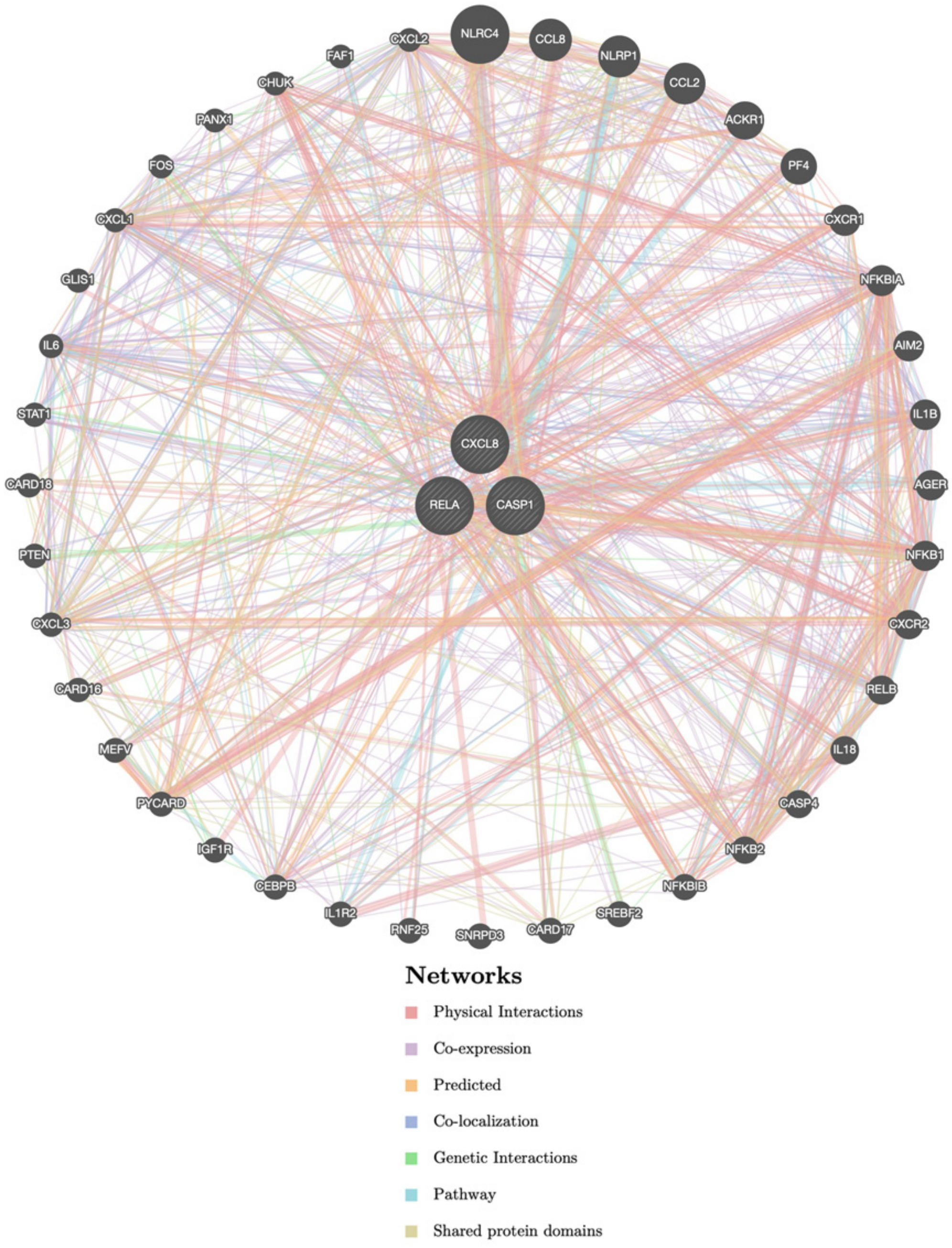


Supplementary Figure 3: Data from the Genemania.org database has shown the interaction of CASP1 with *RelA* and *CXCL-8*.

| **CASP1** | Score | |
|---|---|---|
| Networks | CXCL8 | RelA |
| Physical Interactions | - | 0.031 |
| Co-Expression | 0.014 | 0.007 |
| Predicted | - | - |
| Co-Localization | - | - |
| Genetic Interactions | - | - |
| Pathway | - | - |
| Shared protein domain | - | - |
